# Long-read sequencing identifies GGC repeat expansion in human-specific *NOTCH2NLC* associated with neuronal intranuclear inclusion disease

**DOI:** 10.1101/515635

**Authors:** Jun Sone, Satomi Mitsuhashi, Atsushi Fujita, Takeshi Mizuguchi, Keiko Mori, Haruki Koike, Akihiro Hashiguchi, Hiroshi Takashima, Hiroshi Sugiyama, Yutaka Kohno, Yoshihisa Takiyama, Kengo Maeda, Hiroshi Doi, Shigeru Koyano, Hideyuki Takeuchi, Michi Kawamoto, Nobuo Kohara, Tetsuo Ando, Toshiaki Ieda, Yasushi Kita, Norito Kokubun, Yoshio Tsuboi, Masahisa Katsuno, Yasushi Iwasaki, Mari Yoshida, Fumiaki Tanaka, Ikuo K. Suzuki, Martin C Frith, Naomichi Matsumoto, Gen Sobue

## Abstract

Neuronal intranuclear inclusion disease (NIID) is a progressive neurodegenerative disease characterized by eosinophilic hyaline intranuclear inclusions in neuronal and somatic cells. The wide range of clinical manifestations in NIID makes ante-mortem diagnosis difficult ^1–8^, but skin biopsy realized its ante-mortem diagnosis ^9,10^ and many NIID cases have been diagnosed by skin biopsy^11,12^. Most cases of NIID are sporadic, but several familial cases are known. Using a large NIID family, we conducted linkage mapping, found a 58.1-Mb linked-region at 1p22.1-q21.3 with a maximum logarithm of odds (LOD) score of 4.21, and successfully identified a GGC repeat expansion in the 5’ portion of *NOTCH2NLC* in all affected members by long-read sequencing, but not in unaffected members. We further found the similar expansions in additional 8 unrelated families with NIID as well as 39 sporadic NIID patients. Repeat-primed PCR consistently detected the GGC repeat expansion in all the familial and sporadic patients diagnosed by skin biopsy, but never in unaffected family members nor 200 controls. This shows that pathogenic changes in a human-specific gene evolutionarily generated by segmental duplication indeed causes a human disease.

---

Neuronal intranuclear inclusion disease (NIID), also known as neuronal intranuclear hyaline inclusion disease, or intranuclear inclusion body disease, is a progressive and fatal neurodegenerative disease ^1–3,5^. NIID presents with neuronal and glial intranuclear inclusions widespread through the central and peripheral neuronal system. These inclusions are eosinophilic on hematoxyline and eosin staining, and immunoreactive to anti-ubiquitin and anti-p62 antibodies ^8^. Ultrastructural examination showed that the intranuclear inclusions were well circumscribed masses without a separating membrane composed with mesh of fine straight filaments arranged haphazardly and occasionally with dense core ^3,6,13^. However, neuronal loss is relatively localized in each case, and the extent of neuronal loss is supposed to correlate with the clinical manifestation. Clinical manifestations of NIID are highly variable, and can include pyramidal and extrapyramidal symptoms ^14,15^, cerebellar ataxia ^16,17^, dementia or mental retardation ^1,4^ convulsion ^14^, peripheral neuropathy ^7^, and autonomic dysfunction ^2,7^. This wide range of clinical phenotypes made ante-mortem diagnosis of NIID difficult for a long time. However, by skin biopsy, we have been able to diagnose many NIID cases ^11^. Here we describe a genetic alteration consistently found in *NOTCH2NLC* for familial and sporadic NIID cases.

We examined NIID patients from 9 families (Fig. 1), and 39 sporadic NIID patients. NIID patients of families 1 and 2 showed motor-sensory and autonomic neuropathy ^7^, and patients of families 4, 7, 8, 9, 10 and D showed dementia. NIID patients from family 3 showed both neuropathy and dementia ^11^. Patients of families 1, 2 and 3 presented muscle atrophy and weakness in limbs, limb girdle, and face, and sensory impairment in the distal limbs. Electrophysiologic study showed reduced motor and sensory nerve conduction velocities and amplitudes, and also extensive denervation potentials. In sural nerve specimens, numbers of myelinated and unmyelinated fibers were decreased ^7^.

**Fig. 1.**
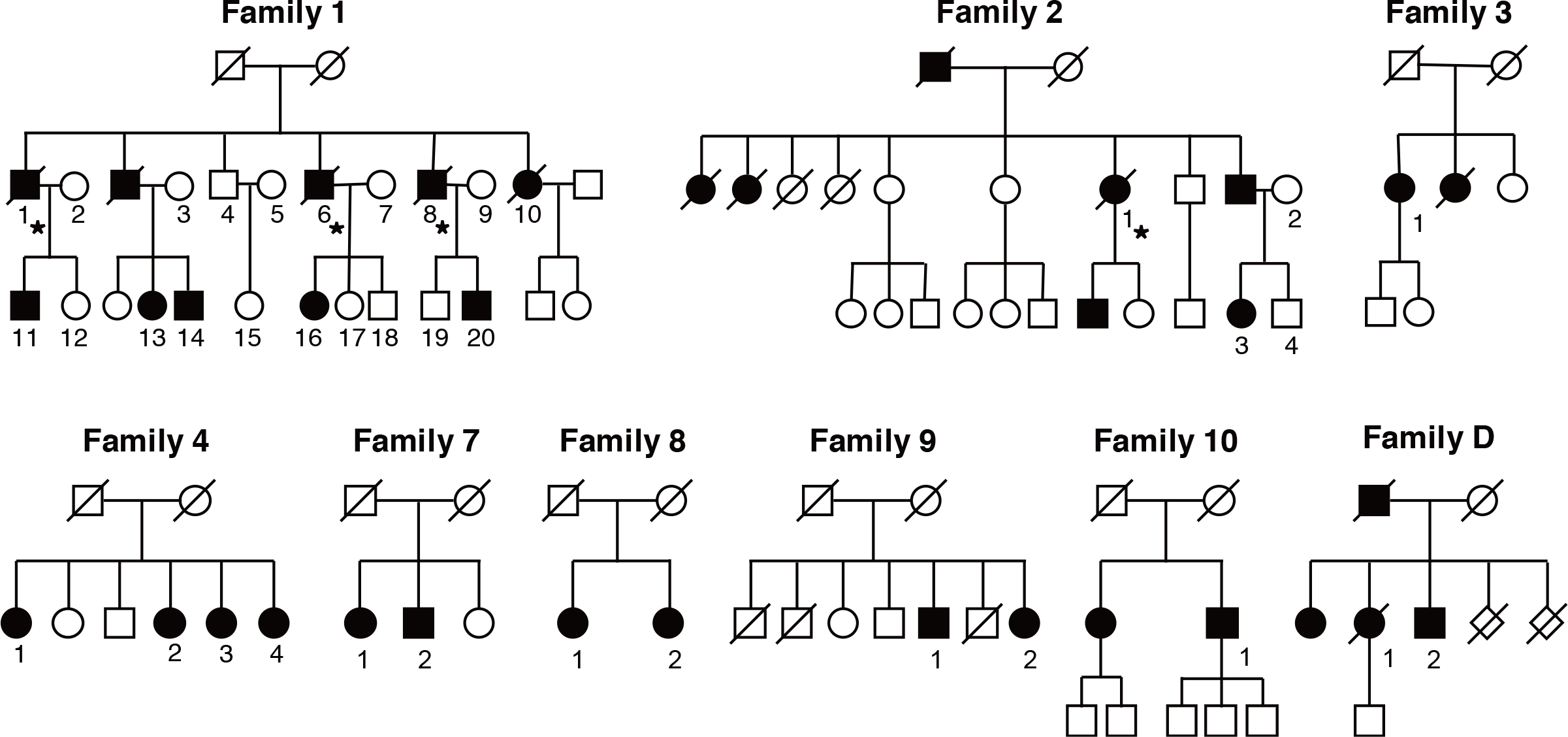
Familial pedigrees. Black filled and white symbols represent affected and unaffected members with familial NIID.

Patients of families 3, 4, 7, 8, 9, 10, D and 39 sporadic NIID cases showed dementia, leukoencephalopathy by T2 weighted image, and high intensity signal in cortico-medurally junction by diffusion weighted image (DWI) in head magnetic resonance imaging (MRI) ^11,18–20^. In autopsies from families 1 and 2 and one sporadic NIID case (3618), eosinophilic, and anti-ubiquitin and anti-p62 positive intranuclear inclusions were observed in central nervous system, peripheral nervous system and visceral organ cells (Fig. 2), and these findings were the basis of NIID diagnosis. Other patients of families 1, 2, 3, 4, 7, 8, 9, 10 and D and other sporadic patients were diagnosed by skin biopsy. All of these familial and sporadic patients were without *FMR1* premutation, the cause of Fragile X associated tremor ataxia syndrome (FXTAS).

**Fig. 2.**
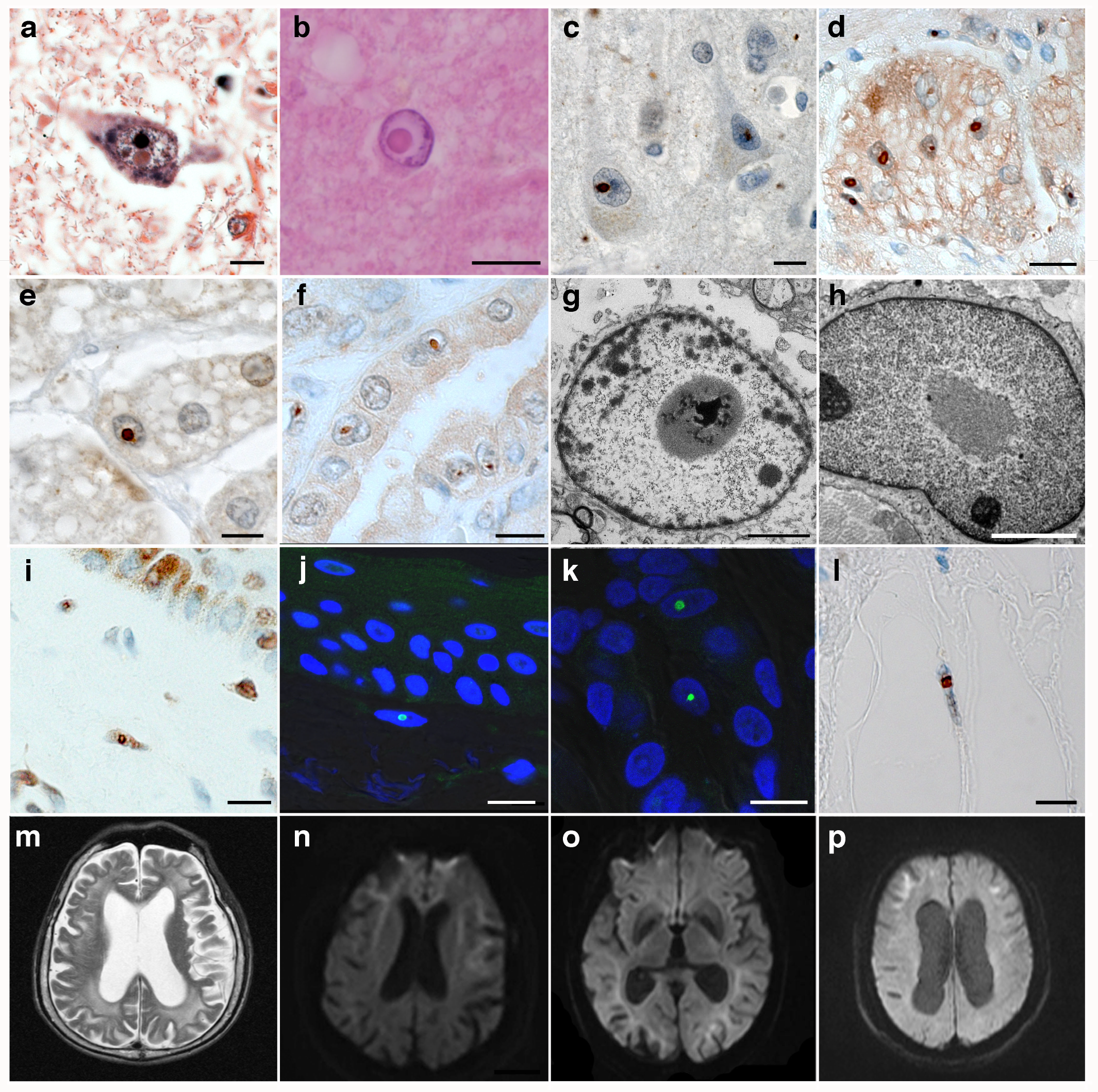
Histopathological features and brain MRI findings of NIID patients. **a-b.** Intranuclear inclusions observed in hematoxylin and eosin stain. **a.** anterior horn neuron in cervical spinal cord (Family 1-8), **b.** astrocyte in cerebral cortex (Sporadic). **c-f.** Immunostaining with anti-ubiquitin antibody. **c.** precentral gyrus (Family1-6), **d.** myenteric plexus (Sporadic), **e.** adrenal cortex (Family 1-1), **f.** renal tubule (Family 1-1). **g-h.** Electron micrograph of an intranuclear inclusion. **g.** astrocyte in cervical spinal cord (Family 1-6), **h.** skin fibroblast (Family1-14). **i-l.** skin biopsy findings with anti-ubiquitin antibody immunostaining. **i**: fibroblast (Family 4-1), **j.** fibroblast (Family 7-2), **k.** sweat gland (Sporadic), **l.** adipocyte (Sporadic). **m-p.** brain MRI findings. **m.** T2 weighted image (Family1-1), **n.** DWI image (Family 3-1), **o.** DWI image (Family 4-2), **p.** DWI image (Sporadic). Scale Bar: 10 μm (**a-f** and **i-l**), 2 μm (**g** and **h**).

First, we performed Illumina Hiseq whole genome sequencing (WGS) or whole exome sequencing (WES) with family 1’s affected and unaffected members. Linkage analysis of family 1 using informative SNPs extracted by LinkDataGen ^21^ from WGS/WES data revealed only two linked-regions with >1.0 maximum logarithm of odds (LOD) scores in chromosome 1 as follows: a 3.5-Mb region at 1p36.31-p36.22 (chr 1: 6218354-9719813 [maximum LOD 2.32]) and a 58.1-Mb region at 1p22.1-q21.3 (chr 1: 94670784-152323132 [maximum LOD 4.21]) based on hg38 (Fig. 3a). We could not identify any pathogenic SNPs or CNVs in WGS or WES data (data not shown).

**Fig. 3.**
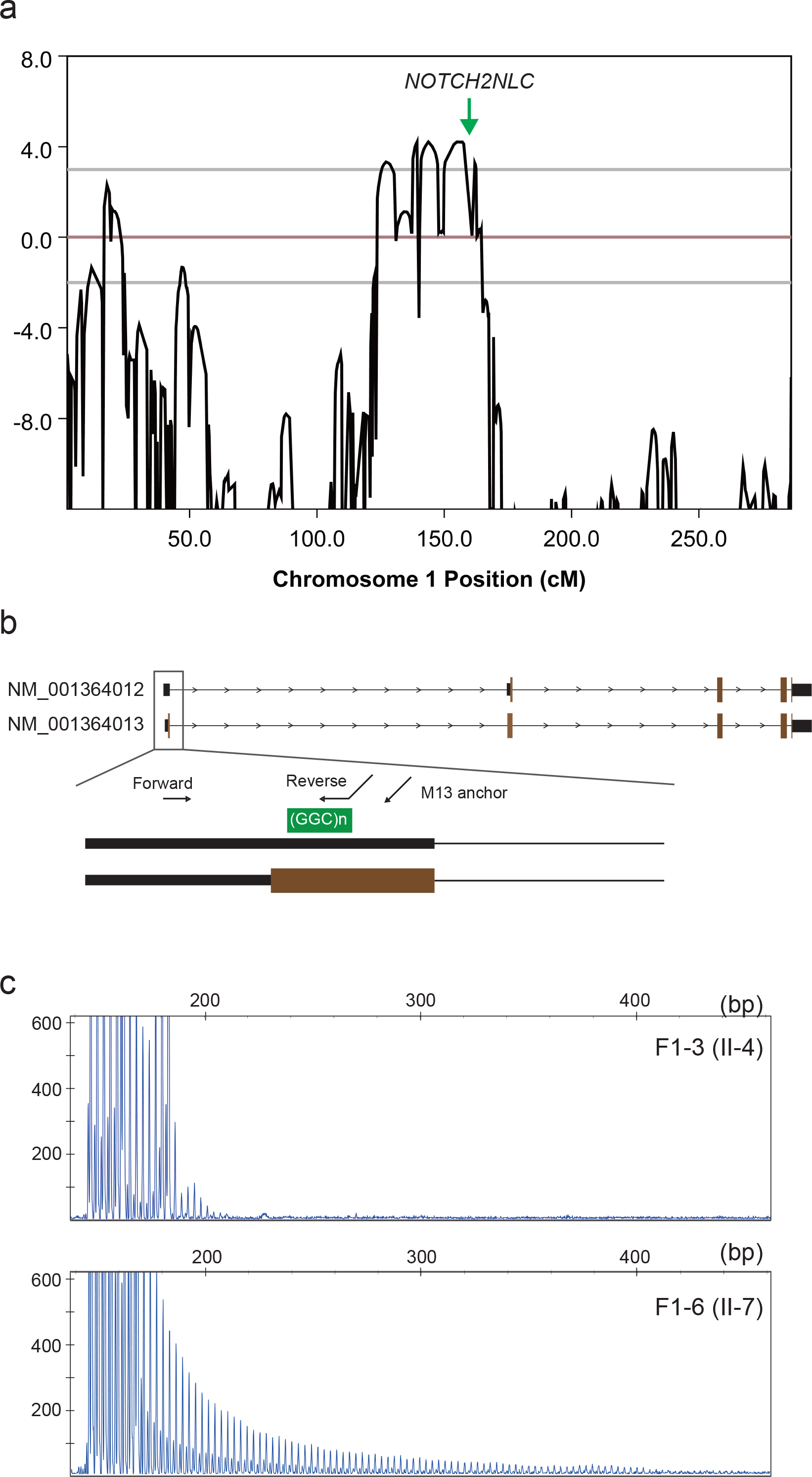
Genetic analysis of family 1. **a.** Linkage analysis identified a 58.1-Mb linked region at 1p22.1-q21.3 with 4.21 maximum logarithm of odds score. **b.** Schematic representation of the *NOTCH2NLC* gene. The disease-associated repeat expansion (green box) was identified in exon 1 of NM_001364012 (isoform 1) and NM_001364013 (isoform 2). Black and brown boxes show untranslated and coding regions, respectively. Three arrows show the primers using repeat-primed PCR. **c.** Electropherogram of repeat-primed PCR. Upper panel shows no repeat expansion in unaffected individual F1-3. Lower panel demonstrates a saw-tooth tail pattern of the repeat expansion in affected individual F1-6.

Then we performed long read WGS either with PacBio RSII or PromethION on 13 affected and 4 unaffected members from 8 families, and checked tandem repeat copy number changes genome-wide using `tandem-genotypes` (https://github.com/mcfrith/tandem-genotypes). Surprisingly, all 13 affected members have GGC repeat expansion at chr1:149390802-149390842, which is in the 5’UTR of the *NOTCH2NLC* gene (NM_001364012) mapped to the 58.1-Mb linked region (Fig. 3b). The repeat expansion was not detected in unaffected family members nor 25 other controls by long read WGS (Figs. 4 and 5, Supplementary Fig. 1 and Supplementary Tables 1 and 2). In all patients, the *NOTCH2NLC* repeat expansions were ranked in the top 5 out of 1,014,212 repeats in the genome (in simpleRepeat.txt from https://www.ucsc.edu) (Supplementary Table 1), using the tandem-genotypes prioritization tool ^22^.

In the reference human genome (hg38), this repeat sequence is (GGC)_9_(GGA)_2_(GGC)_2_. It is possible that expanded repeats contain GGA in addition to GGC. Expanded repeat sequences were extracted and multiple-alignment was done using MAFFT (Supplementary Fig. 2) ^23^. Due to nanopore’s high tendency of errors in homopolymers and adenine/guanine substitutions, it is difficult to obtain accurate multiple alignment and consensus sequence of expanded repeats. However, it is possible that (GGA)n is also present in the expanded repeat towards the 3’ end in the patients from families 1, 2 and 3, because reads from both DNA strands support GGA, and also PacBio reads in one individual (F1-6) have GGA (Supplementary Fig. 2, asterisk).

**Fig. 4.**
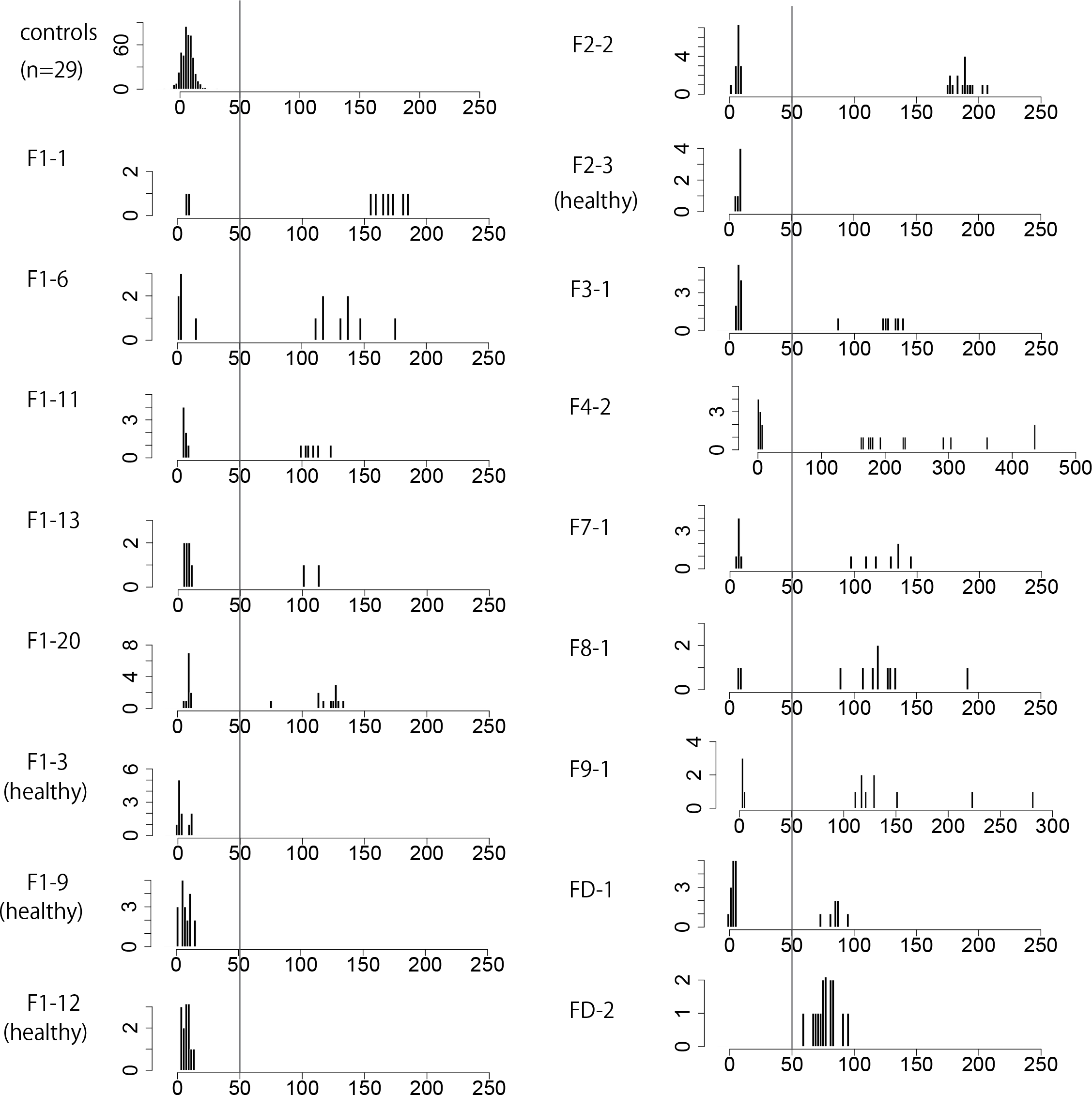
Expanded repeats detected by tandem-genotypes. tandem-genotypes prediction shows that all affected individuals from 8 families have expanded *NOTCH2NLC* repeat. The *NOTCH2NLC* repeat is at chr1:149390802-149390842, containing (GGC)_9_(GGA)_2_(GGC)_2_G according to simpleRepeat.txt from UCSC. y-axis: read count, x-axis: change in copy number relative to the reference human genome. The +50 copy number changes is marked by a line: none of the control and unaffected individuals has repeat expansion beyond this line. Repeat copy number changes from the refernece in controls ranges from −13 to 32 (median: 7), and that of NIID has bimodal distribution (predicted normal allele: 0-16 (median: 7) and predicted expanded allele: 60-436 (median: 128)). Note the x-axis ranges in F4-2 and F9-1 are different from others. For controls, combined repeat copy number changes from 29 non-NIID controls are plotted in the same histogram (see Supplementary Fig. 1).

**Fig. 5.**
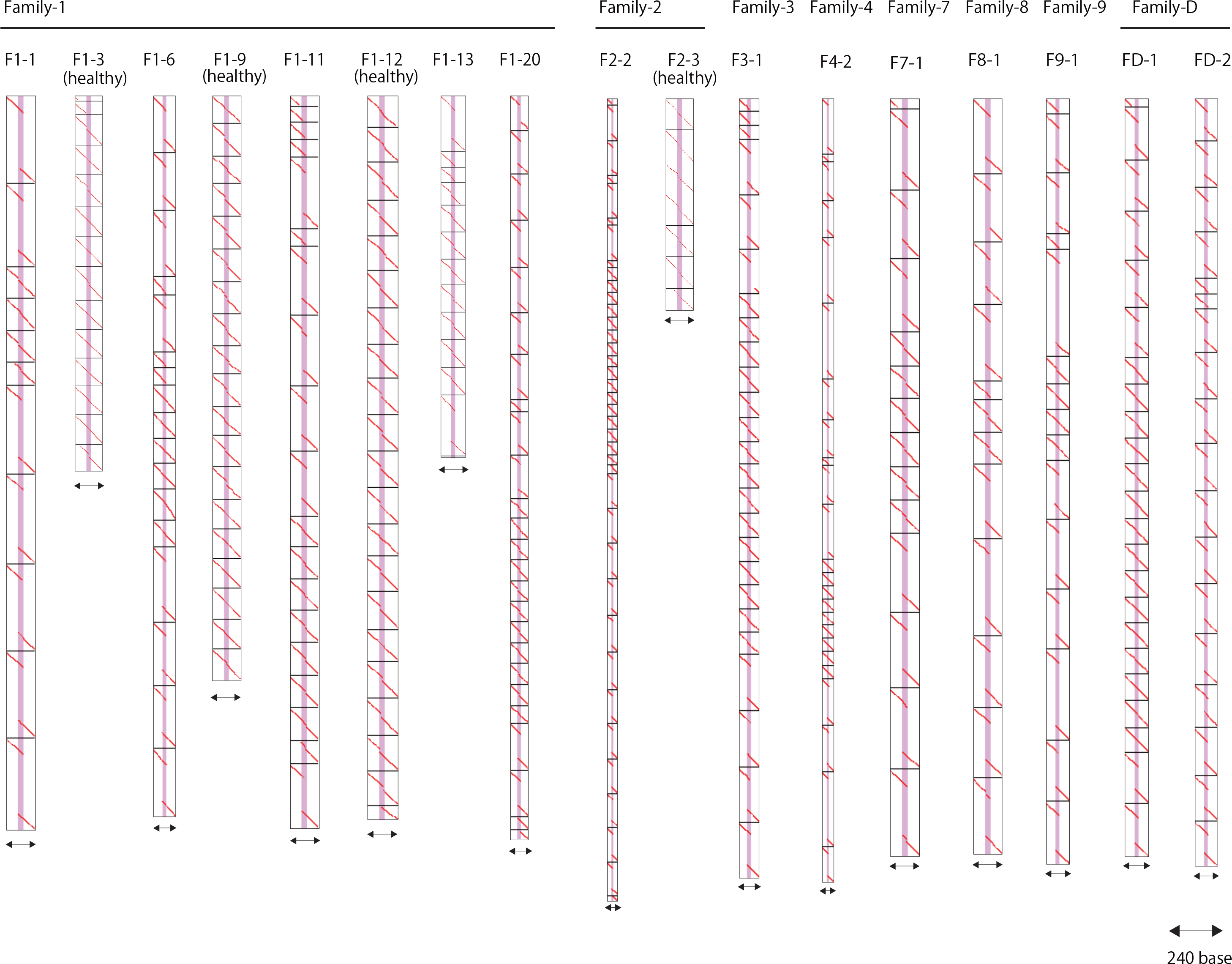
Dot-plot presentation of expanded and non-expanded repeats in familial NIID patients. Dot-plot pictures of *NOTCH2NLC* repeat expansions from all members of NIID families, showing alignments of DNA reads (vertical) to the reference human genome (horizontal). Horizontal two-headed arrow shows the length of the reference (240 bases in all dot-plots). The vertical pink-gray line represents the *NOTCH2NLC* repeat (chr1: 149390802-149390842).

Next, we investigated the segregation of this repeat expansion in genomes of other patients in 9 families, and unaffected members of families 1 and 2, with repeat-primed PCR (RP-PCR). RP-PCR demonstrated the repeat expansion as a saw-tooth pattern in all the tested patients with NIID but not in unaffected familial members nor in 200 normal controls (Fig. 3c and Supplementary Table 3). We also checked the repeat expansion in 39 sporadic NIID patients. Surprisingly, all the sporadic patients had the expansion (Supplementary Table 4). Taken together, (GGC)n repeat expansion in *NOTCH2NLC* was consistently found in both familial NIID patients and sporadic NIID patients diagnosed by skin biopsy.

It is not clear whether expanded GGC repeats become highly methylated, which might possibly repress *NOTCH2NLC* transcripts from the expansion allele. The 5-methylcytosine (5mC) modification in both expanded and non-expanded alleles was examined from nanopore sequencing signal data, using a recently available nanopore basecaller, `flappie vl.1.0` (https://github.com/nanoporetech/flappie) (Supplemental methods). There was no difference in the rate of 5mC between expanded and non-expanded repeats (Supplementary Fig. 3). Comparing 5mC within the repeat may be difficult because the repeat sequences of expanded and non-expanded alleles are different. So we also examined the CpG islands (chr1:149390845-149391541) immediately downstream of the repeat and compared the number of 5mC on the expanded and non-expanded repeat alleles. There is no difference between the alleles (Supplementary Fig. 4). Our 5mC measurement did not support enhanced DNA methylation of the 5’UTR and intron 1 of *NOTCH2NLC* in patients, however, it does not exclude a possibility of allelic repression due to mRNA instability or reduced efficiency of translation due to repeat expansion in the 5’UTR of *NOTCH2NLC*.

*NOTCH2NLC* is one of the three human-specific *NOTCH2*-related genes (*NOTCH2NLA*, *NOTCH2NLB* and *NOTCH2NLC*) in 1q21.1, highly expressed in various radial glia populations including outer radial glia, astrocytes and microglia, and thought to be involved in the evolutionary expansion of the human cortex ^24,25^ The *NOTCH2NL* genes have >99.1% sequence identity to the N-terminal portion of *NOTCH2*, but *NOTCH2NLC* contains a 2 bp deletion just downstream of the *NOTCH2* start codon and lacks the N-terminal signal peptide. Ectopic expression of *NOTCH2NL* delays cortical neuron differentiation in mouse, and deletion of *NOTCH2NL* affects human cortical organoid development ^24^. The onset ages of NIID patients in this study are after 20 years, no patients showed structural brain abnormalities such as microcephaly, and DNA methylation of *NOTCH2NLC* showed no difference between GGC expanded and non-expanded alleles. Therefore, the functional loss of *NOTCH2NLC* through CpG methylation due to GGC expansion may be less likely.

In this study, NIID patients showed (GGC)n expansion in the 5’UTR of *NOTCH2NLC*, and autopsied NIID cases consistently present with ubiquitin-positive intranuclear inclusions in both nervous system and somatic cells (when tested). The combination of triplet repeat expansion and widely observed intranuclear inclusion is similar to Fragile X-associated tremor/ataxia syndrome (FXTAS) ^12,26^. Expansion of a (GGC)n repeat to 55-200 copies in the 5’UTR of *FMR1* causes FXTAS, presenting tremor, ataxia and cognitive dysfunction in the sixth decades with widely observed intranuclear inclusions both in nervous system cells and somatic cells ^27,28^. One hypothesis of the FXTAS toxicity mechanism is that repeat-bearing RNA interacts with multiple GGC RNA-binding proteins leading to their functional depletion ^29^. Another hypothesis is that GGC repeats in the Fragile X RNA can be translated into toxic proteins, either by repeat-associated non-AUG (RAN) translation or the utilization of noncanonical start codons to produce expanded polypeptides, such as polyglycine ^28,30,31^. As for NIID, both of these hypotheses may possibly be related to the pathomechanism of NIID, and we need to investigate the expression or translation of GGC expanded RNA, including the constituents of intranuclear inclusions, in the next step. Besides, in this study, families 1 and 2 showed weakness, families 4, 7, 8, 9, 10 and D showed dementia and leukoencephalopathy, and families 3 showed both weakness and dementia. We could not recognize any difference or trend in the degree of (GGC)n expansion among these three subgroups, perhaps due to inaccuracy of long read sequencing data. Furthermore, we observed similar (GGC)n expansion in all 39 sporadic NIID patients. The clinical manifestation of these sporadic NIID patients was almost the same as the familial NIID patients with dementia, the onset age was around the sixties and head MRI showed leukoencephalopathy and abnormally high intensity DWI signal in the cortico-medullary junction ^11^. Therefore, except for familial presentation, we can not practically differentiate familial and sporadic NIID. We need to study on these unsolved issues of NIID with a large NIID patient cohort diagnosed by skin biopsy and genetic alteration in *NOTCH2NLC*, and hopefully develop a disease modifying therapy of NIID in the near future.

The *NOTCH2NL* genes at 1q21.1 were not annotated as three different genes, because these regions were erroneously assembled and missed in the human reference genome, until hg38 ^24,25^. These regions are highly similar, moreover (GGC)n is 100% GC-rich, which is difficult to analyze by standard sequencing methods. Long read sequencing with >10 kb reads may have a big advantage for resolving such homologous (duplicated) and repetitive regions, though its application to actual human diseases is still challenging ^32^. We took advantage of using PromethION and applied it to multiple patients, and obtained X8~X25 whole genome coverage from single flowcell runs. In addition, we recently developed a computational tool, tandem-genotypes ^22^, to find tandem repeats possibly causing human diseases. By utilizing this, we detected *NOTCH2NLC* repeat expansions in familial and sporadic NIID, even from as low as (X8) coverage data. Therefore, this study provides a stark example of long read sequencing for resolving unsolved human diseases.

To our knowledge, clear pathogenic variants have never been reported among approximately 30 human-specific genes evolutionarily generated by segmental duplications ^24,25,33^. Thus, we believe that such human-specific genes are very important not only in human (brain) evolution but also in human neurological diseases. We anticipate that further studies will reveal additional pathogenic variations in human specific genes associated with human diseases.

In conclusion, the genetic cause of familial and sporadic NIID is finally solved. We predict that the prevalence of NIID is larger than previously thought ^11^, and NIID may possibly account for a substantial part of dementia population. To date, many NIID cases remain undiagnosed with different clinical diagnosis such as Binswanger’s disease or neurodegenerative disease with white matter damage. NIID should be considered as a differential diagnosis for dementia, and its genetic diagnosis has been done histopathologically and can be now performed genetically. It may be worth investigating the *NOTCH2NLC* repeat expansions in a large population with dementia.

## URLs

**LinkDataGen**: http://bioinf.wehi.edu.au/software/linkdatagen/

**Merlin**: http://csg.sph.umich.edu/abecasis/merlin/

**LAST**: http://last.cbrc.jp

**tandem-genotypes**: https://github.com/mcfrith/tandem-genotypes

**MAFFT**: https://mafft.cbrc.jp

## Acknowledgements

We thank all the patients and their families for participating in this study. We also thank Drs. Osamu Komure, Naoyuki Kitagawa, Hajime Yoshimura, Junko Ishii, Kyoko Higashida, Masaya Togo, Tatsuhiko Yuasa, Hiroyuki Nakayasu, Yutaka Suto, Tateo Manabe, Makoto Takahashi, Mai Tsutiya, Naoko Uehara, Houdou Mori, Takashi Tokunaga, Takashi Inuzuka, Akira Takekoshi, Shota Anzai, Kimito Kondo, Tetsuya Takahashi, Kazuki Muguruma, Yoshiko Sugihara, Ken Yokote, Shogo Takamura, Nobuki Oohara, Eri Hayano, Keiko Saiki, Daiji Hori, Yuishin Izumi, Rei Kobayashi, Mika Saiki, Yuka Tsukahara, Masaru Kuriyama, Takashi Kurashige, Yoshiaki Takahashi, Tomoko Noda, Shinnosuke Takagi, Kazuhiro Honda, Masumi Ito, Aya Yarita, Yuki Satake, Tomonori Inagaki, Keita Hiraga, Yoshiyasy Kato and many neurologists for evaluating clinical manifestation in NIID patients and supporting the diagnosis of NIID patients. This work was supported by AMED under grant numbers, JP18ek0109280, JP18dm0107090, JP18ek0109301, JP18kk0205001, JP18ek0109348, JP18md0107059, JP18ek0109284, and JP18dm020715; JSPS KAKENHI Grant Numbers, 17K15630, JP17K16132, 17H06994, 24591257 and 15K09312; Takeda Science Foundation; Daiwa Securities Health Foundation;Termo Foundation for Life Sciences and Arts.

## Author contributions

J.S., A.H., H.T., Y.K., H.S., Y.Ta, K.M., T.A., T.I., Y.K., N.K., Y.Tsu, H.D., S.K., H.T., H.K., M.K., F.T. and G.S. access the clinical manifestation of NIID patients of this study. J.S., K.M., H.K., Y.I., and M.Y. performed the histopathological experiments and interpreted their data. J.S., S.M., A.F., T.M., M.C.F., N.M. and G.S. performed the genetic experiments and interpreted their data. J.S., S.M., A.F., N.M., and G.S. wrote the manuscript with contribution from all remaining authors.

