## Supplementary material for "Long-read sequencing identifies GGC repeat expansion in human-specific *NOTCH2NLC* associated with neuronal intranuclear inclusion disease"

17. Department of Neuropathology, Institute for Medical Sciences of Aging, Aichi Medical University, Nagakute, Japan.
18. Universite Libre de Bruxelles, Institute for Interdisciplinary Research and ULB Neuroscience Institute, Brussels, Belgium.
19. Artificial Intelligence Research Center, National Institute of Advanced Industrial Science and Technology, Tokyo, Japan.
20. Graduate School of Frontier Sciences, University of Tokyo, Chiba, Japan.
21. Computational Bio Big-Data Open Innovation Laboratory, National Institute of Advanced Industrial Science and Technology, Tokyo, Japan.
22. Department of Neurology, and Brain and Mind Research Center, Nagoya University Graduate School of Medicine, Nagoya, Japan.
23. These authors contributed equally: Jun Sone, Satomi Mitsuhashi, Atsushi Fujita
24. These authors jointly supervised this work: Naomichi Matsumoto, Gen Sobue
*

### Supplementary Methods

#### Samples of NIID families

Genomic DNA of peripheral blood leukocytes was extracted from affected and non-affected members of 9 NIID families and 39 sporadic NIID patients after obtaining informed consent. All the affected patients were diagnosed by skin biopsy. Study protocols were approved by institutional review boards of Yokohama City University Faculty of Medicine and Nagoya University School of Medicine.

#### Pathology

Specimens were fixed in 20% formaldehyde and embedded in paraffin. They were cut into 6-μm slices and stained by the hematoxylin and eosin. Immunistaining and immunofluorescence staining were performed as described previously [1] with anti-ubiquitin antibody (Z0458, Dako) and Ventana Discovery system (Roche, Basel, Schweiz), using the Ventana DAB map kit (Roche, Basel, Schweiz). Samples for electron microscopic study were prepared as described previously [2].

#### *FMR1* gene analysis

Genomic DNA was extracted from peripheral blood samples of these patients with informed consent. Southern blotting was performed as described previously [3]. We defined normal alleles as range of 5 to 44 repeats, and premutation alleles as 55 to 230 repeats.

#### Linkage analysis

We performed linkage analysis for family 1 using whole exome sequencing (F1-4, 10, 11, 12, 13, 14, 19, 20 and 24) or whole genome sequencing (F1-6, 7, 16, 17 and 18) data of seven affected and seven unaffected individuals. The informative genotyping data was selected by LinkDataGen [4] and linkage analysis was performed with Merlin [5] using the condition of the autosomal dominant inheritance with complete penetrance.

#### Long read whole genome sequencing of NIID patients

DNA samples of human patients with NIID phenotype and unaffected family members were sequenced by PromethION nanopore sequencer (Oxford Nanopore Technologies). Library preparation was done using 1D Genomic DNA ligation kit (SQK-LSK109) according to manufacturer's protocol. For each individual, one PRO-002 (R9.4.1) flowcell was used. Base-calling and fastq conversion were performed with MinKNOW (v1.14.0) (Oxford Nanopore Technologies). One individual sample

(F1-6) was sequenced by RSII sequencer (PacBio). Control datasets were either sequenced by MinION (Oxford Nanopore Technologies), PromethION or Sequel (PacBio).

**Long read sequencing datasets from control individuals**

Long read whole genome sequencing datasets of non-NIID individuals were obtained to investigate genetic alterations in patients with other diseases or their family members, either using MinION/PromethION or Sequel sequencers. Publicly available human whole genome nanopore (rel3) sequence data from one individual (NA12878) were also used [6]. Another publicly available 60X coverage human whole genome nanopore dataset from a different individual (NA19240) using PromethION was downloaded from https://www.ebi.ac.uk/ena/data/view/PRJEB26791 [7]. For MinION sequencing, library preparation was done using a 1D genomic DNA kit (SQK-LSK108) and sequenced with FLA-MIN106 (R9.4.1) flowcells, according the manufacturer's protocol. Base-calling and fastq conversion were performed with MinKNOW ver1.11.5.

**Long read sequence data analysis using LAST**

Reads were aligned to human reference genome (hg38) using LAST (http://last.cbrc.jp) version 936 or 959 as follows [8]:

```
windowmasker -mk_counts -in hg38.fa > genome.wmstat
windowmasker -ustat genome.wmstat -outfmt fasta -in genome.fa > genome-wm.fa
lastdb -P8 -uNEAR -R11 -c GRCh38 genome-wm.fa
last-train -P8 GRCh38 reads.fasta > train.out
lastal -P8 -p train.out GRCh38 reads.fasta | last-split > alns.maf
```

Then tandem repeat genotyping and multi-dataset prioritization were done using tandem-genotypes v1.1.0 (https://github.com/mcfrith/tandem-genotypes, also in bioconda) as described elsewhere [9]. Tandem repeat (simpleRepeat.txt) and gene (refFlat.txt) annotations were obtained from the UCSC genome database (http://genome.ucsc.edu/) [10]. For all NIID patients, two publicly available human datasets (NA12878 nanopore rel3 and NA19240 PromethION ERR2585112-5) and also two unaffected family members (F1-3 and F1-9) were used to de-prioritize possibly-benign expansions.

The actual commands used were:

```
tandem-genotypes -g refFlat.txt simpleRepeat.txt alns.maf > patient-out
```

```
tandem-genotypes-join patient-out : controls-out > patient-prioritized
```

Dot-plot pictures for the *NOTCH2NLC* repeat were obtained as follows:

```
last-dotplot -1 chr1:149390802-149390842 --sort2=3 –strands2=1 –rot1=v –rot2=h –labels1=2 –max-gap2=0,inf –bed1 notch2nlc-repeat.bed alignment.maf alignment.png
```

**Extracting repeats from long reads**

Repeat sequences aligned to chr1:149390802-149390842 were extracted with flanking 50-bp sequences, then multiple-alignment was done using MAFFT version 7 [11].

**Repeat-primed PCR**

Genomic DNA of affected 9 families and 39 sporadic patients and 200 normal control individuals were analyzed by repeat-primed PCR (RP-PCR). The PCR mix contained 0.25U PrimeSTAR GXL DNA Polymerase, 1×PrimeSTAR GXL Buffer, 200 μM each dATP, dTTP, dCTP (Takara Bio, Shiga) and 7-Deaza-2′-deoxy-guanosine-5′-triphosphate (Sigma-Aldrich, St Louis, MO, USA), 5% dimethyl sulfoxide (Sigma-Aldrich), 1M betaine (Sigma-Aldrich), 0.3 μM each primer mix and 100 ng genomic DNA in a total reaction volume of 10 μl. The primer mix contained three primers, NOTCH2NLC-F; 5'-FAM-GGCATTTGCGCCTGTGCTTCGGACCGT-3', M13-$(GGC)^4(GGA)^2$-R; 5'- CAGGAAACAGCTATGACCTCCTCCGCCGCCGCCGCC-3', and M13-linker-R; 5'-CAGGAAACAGCTATGACC-3'. After incubation at 98 ℃ for 10 minutes, the cycling conditions were followed by 16 cycles of 98 ℃ for 30 seconds, 66 ℃ for 1 minute with reduced 0.5 ℃ per cycle and 68 ℃ for 8 minutes, followed by 29 cycles of 98 ℃ for 30 seconds, 58 ℃ for 1minute and 68 ℃ for 8 minutes, followed by final elongation step of 68 ℃ for 10 minutes. The ramp rate of all cycling step was adjusted 0.5℃ per second. Electrophoresis was performed on a 3500xl Genetic analyzer (Thermo Fisher Scientific, Waltham, MA, USA) and the data was analyzed using GeneMapper software (Thermo Fisher Scientific). We judged a saw-tooth tail pattern in the electropherogram as the disease-associated repeat expansion.

**Nanopore 5mC methylation modification calling**

We called 5mC methylation using a nanopore basecaller, flappie v1.1.0 (https://github.com/nanoporetech/flappie) like this:

```
flappie --model r941_5mC fast5 > fastq
```

We tested two *NOTCH2NLC* regions in affected NIID patients; the expanded repeat itself and the downstream CpG island, chr1:149390802-149390842 and chr1:149390845-149391541, respectively. We also tested the G-quadruplex

(CGGGGG) on chr15:24848213-24848273, which is in a CpG island of *SNRPN* [12]. This region is maternally imprinted and known to be responsible for Prader-Willi syndrome. (We expect to see a bimodal methylation pattern here.)

In the obtained sequences, flappie describes 5mC as “Z”. We aligned these sequence to human reference genome hg38 as described above. Reads aligned to the G-quadruplex of *SNRPN* or the $(GGC)_n$ repeat of *NOTCH2NLC* were extracted, and the number of “Z” per 1,000 bases was shown in a histogram (Supplementary Figs. 3 and 4).

### Supplementary figures

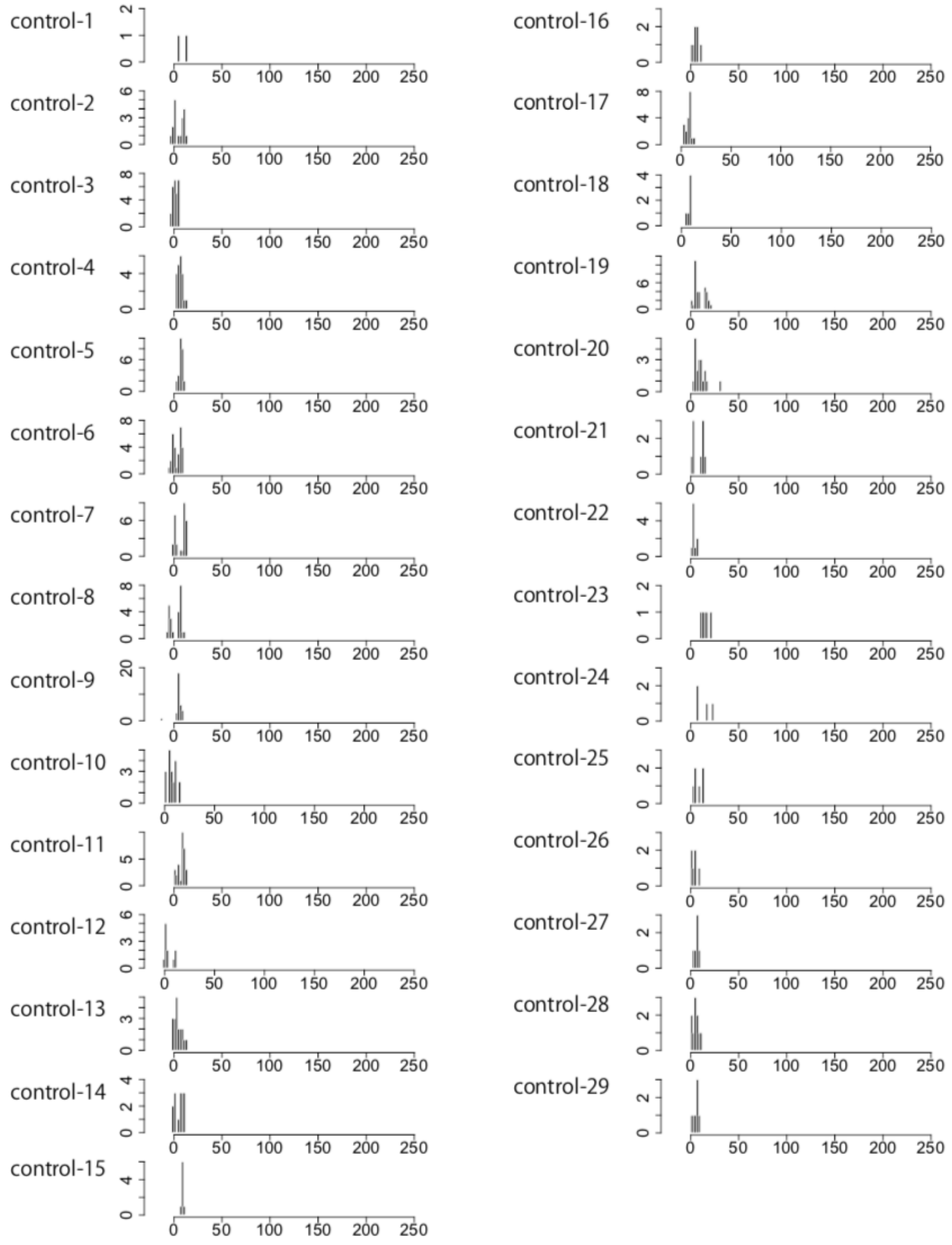


**Supplementary Fig. 1 GGC repeats in normal controls**

`tandem-genotypes` prediction of normal controls in *NOTCH2NLC* repeat. y-axis: read count, x-axis: change in repeat copy number relative to the reference human genome.

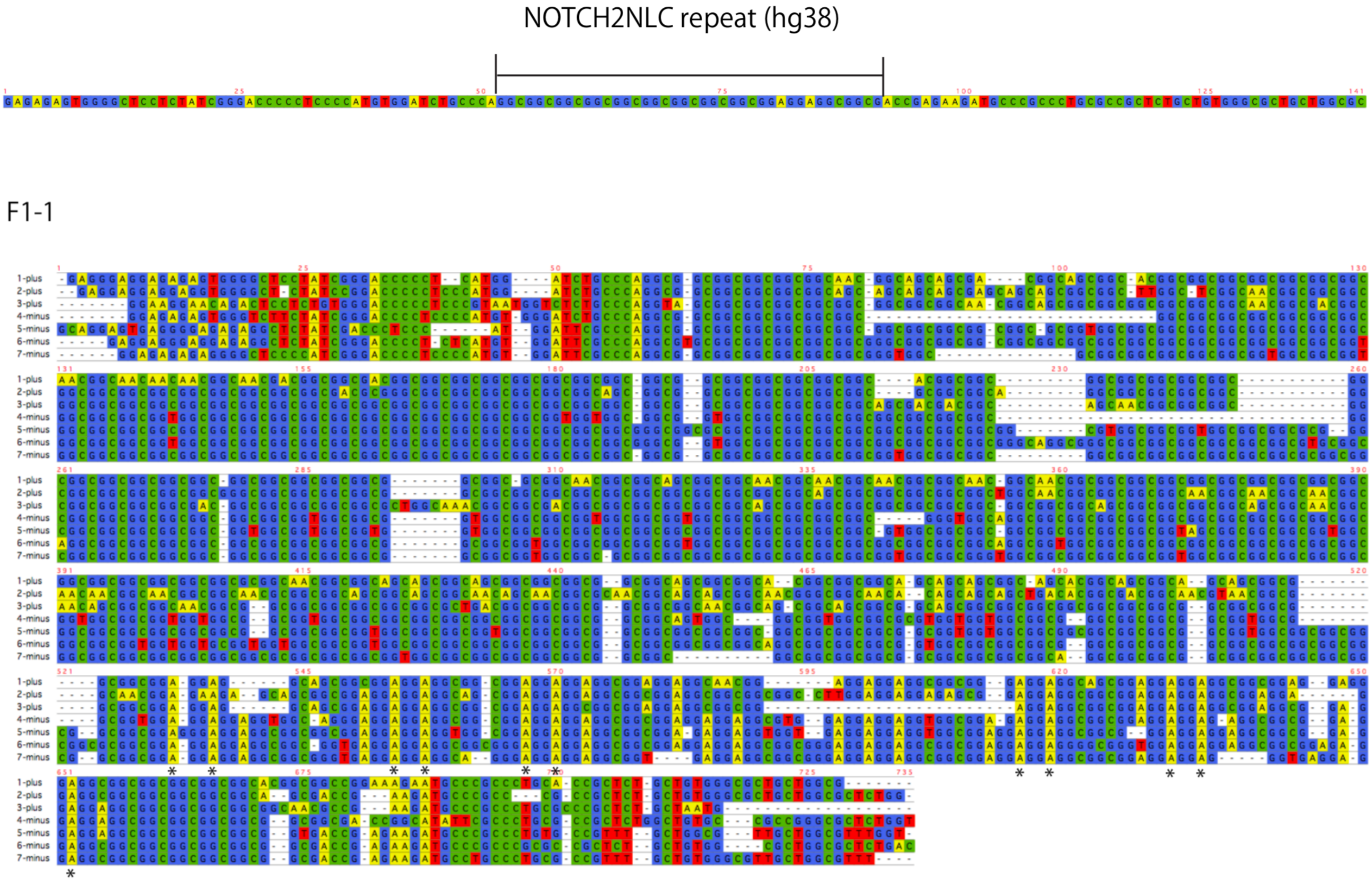
NOTCH2NLC repeat (hg38)
F1-1
1-plus
2-plus
3-plus
4-minus
5-minus
6-minus
7-minus

F1-6 (PacBio reads)

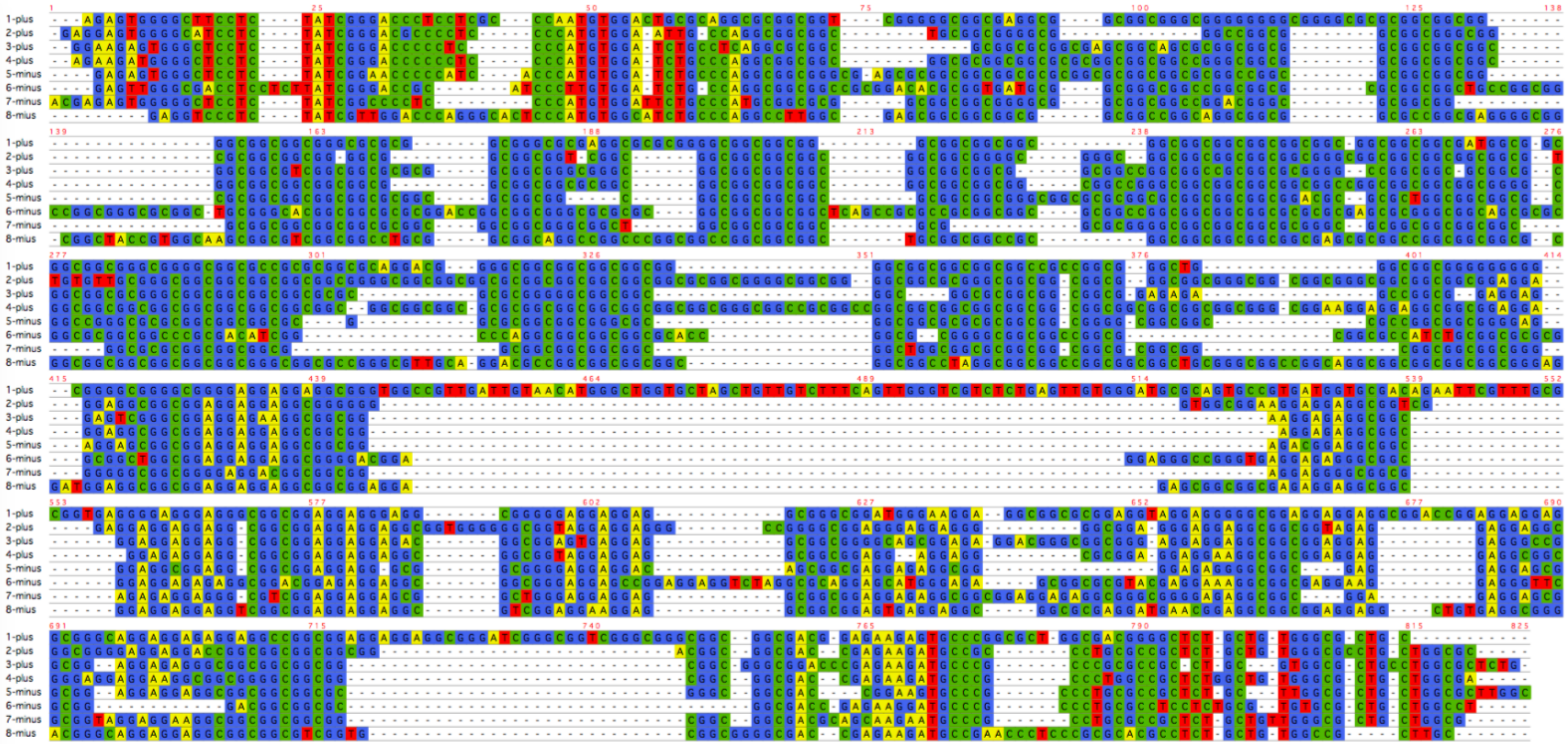

F1-11

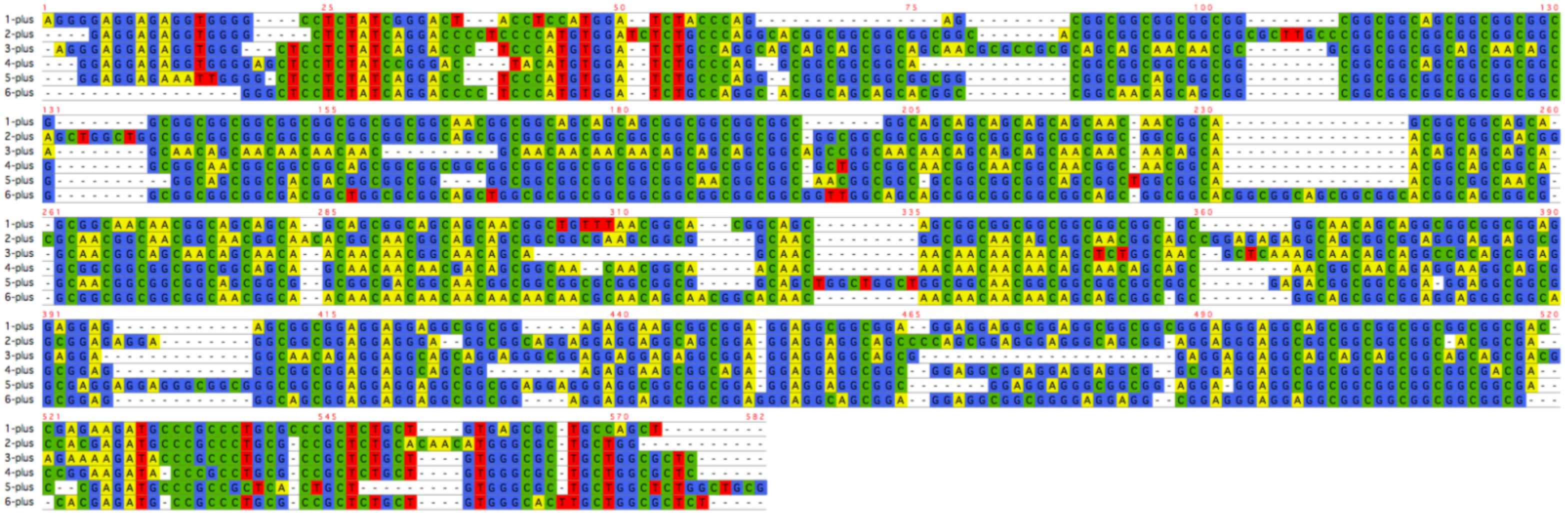

F1-13

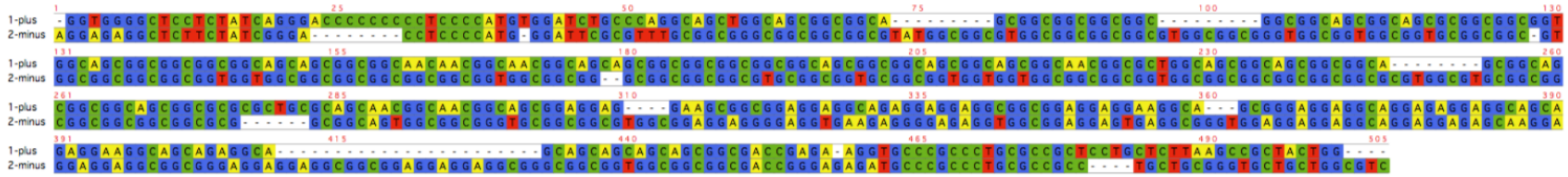

F1-20

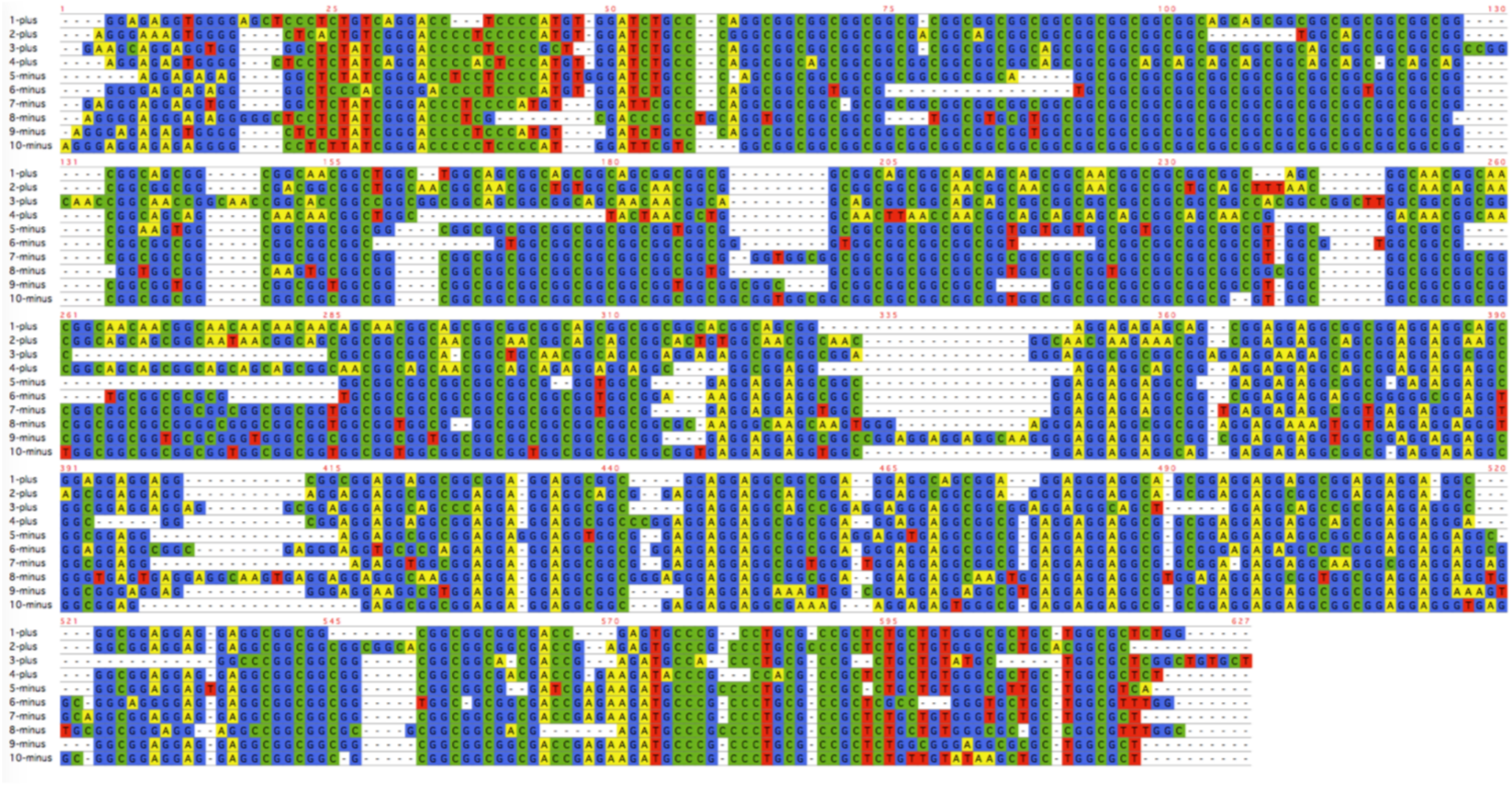

# F2-2

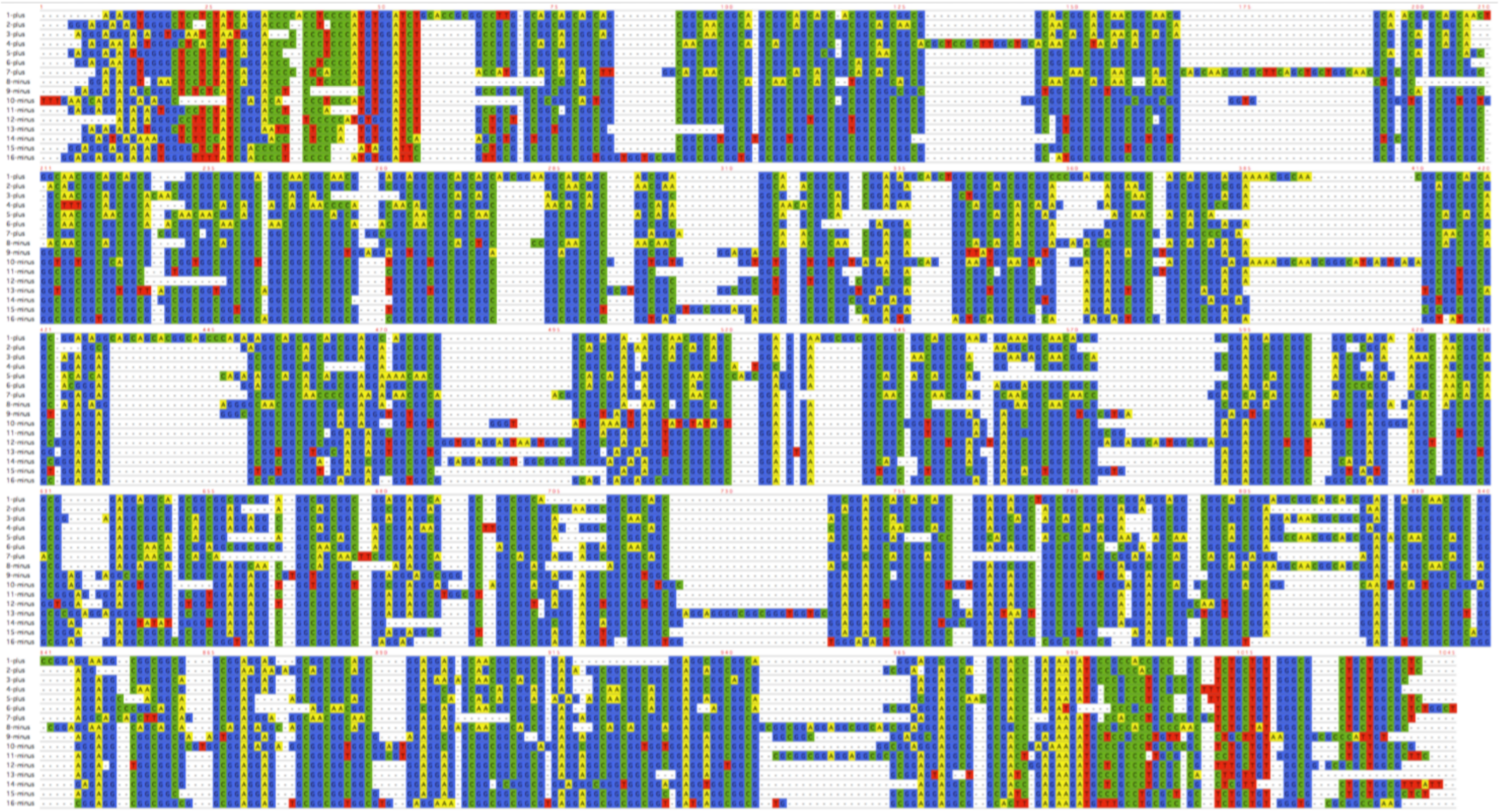

F3-1

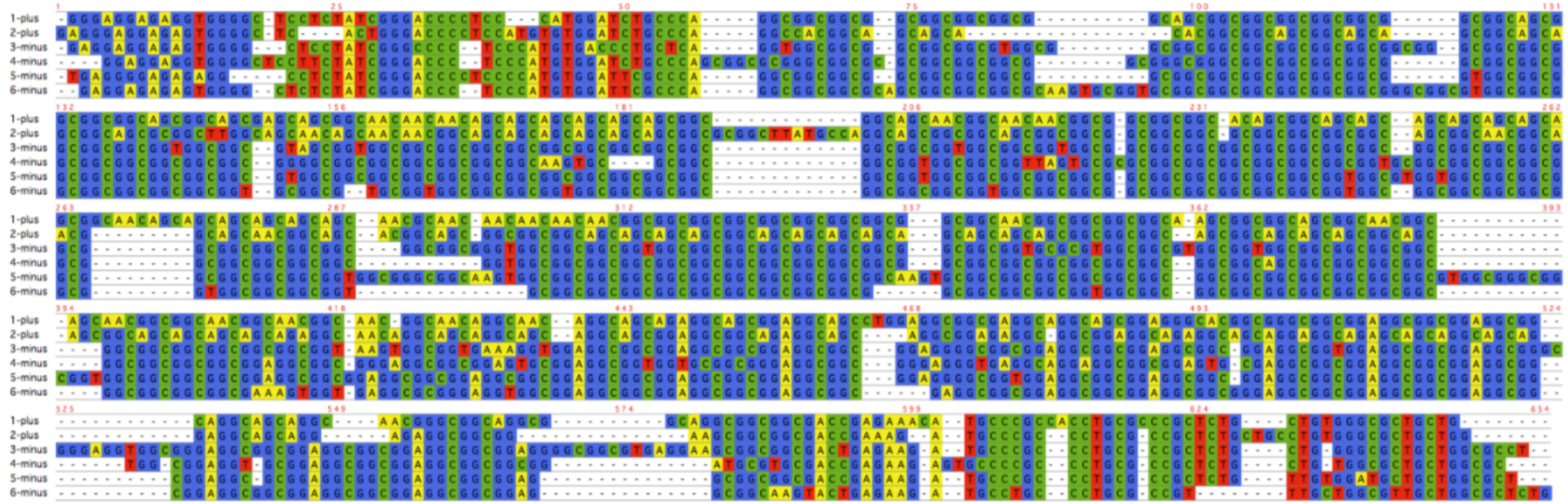
1-plus
2-plus
3-minus
4-minus
5-minus
6-minus

F7-1

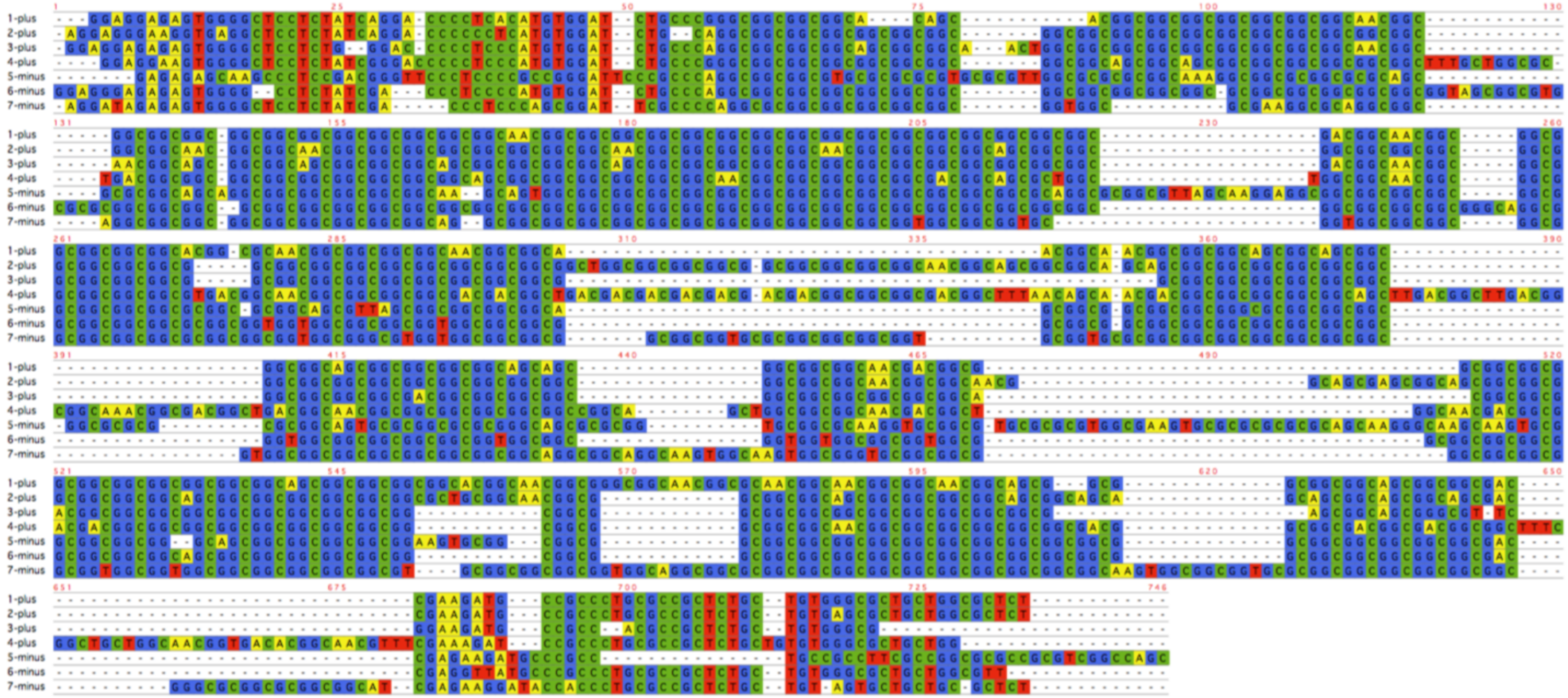
1-plus
2-plus
3-plus
4-plus
5-minus
6-minus
7-minus

F4-2

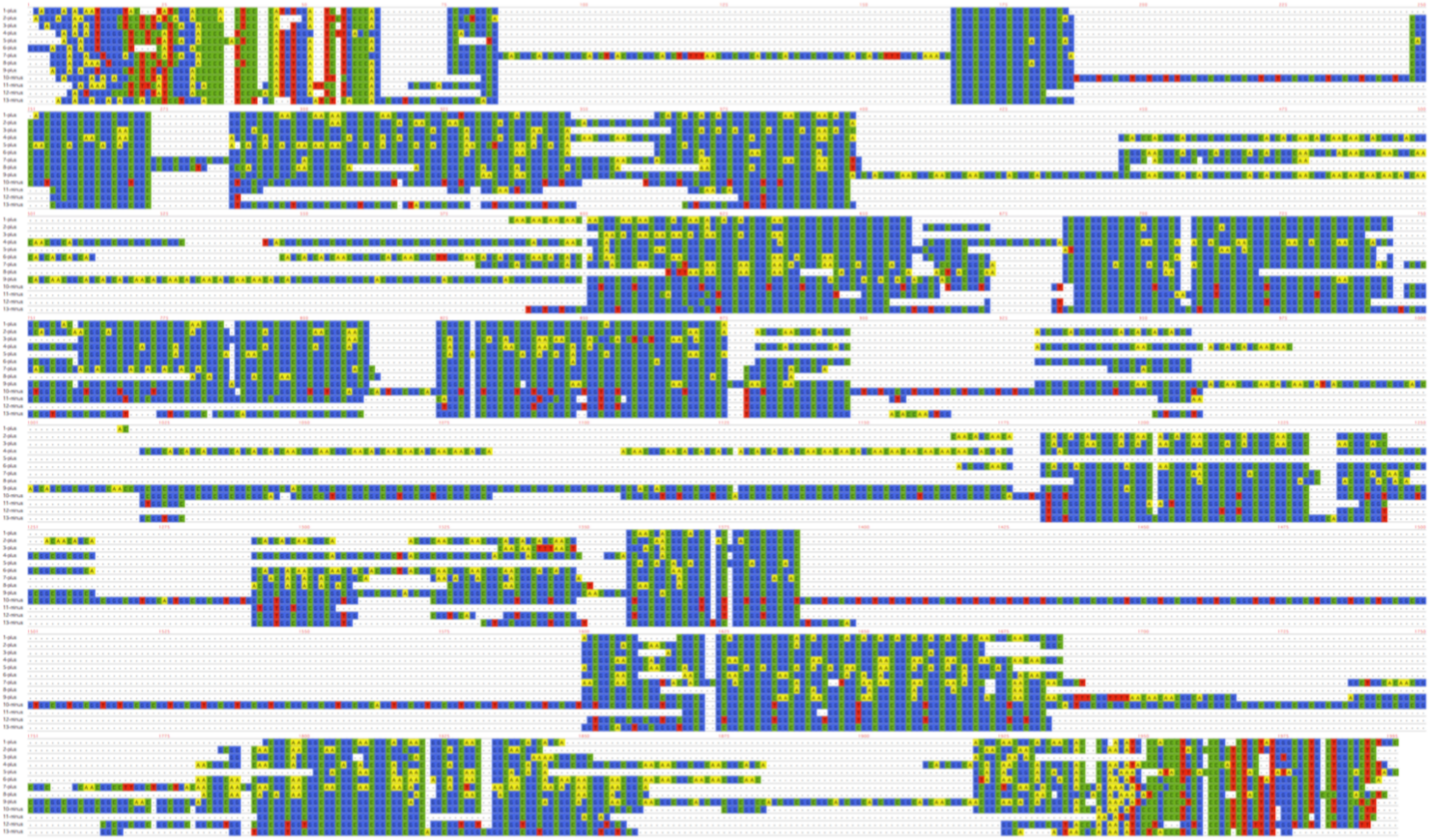

F8-1

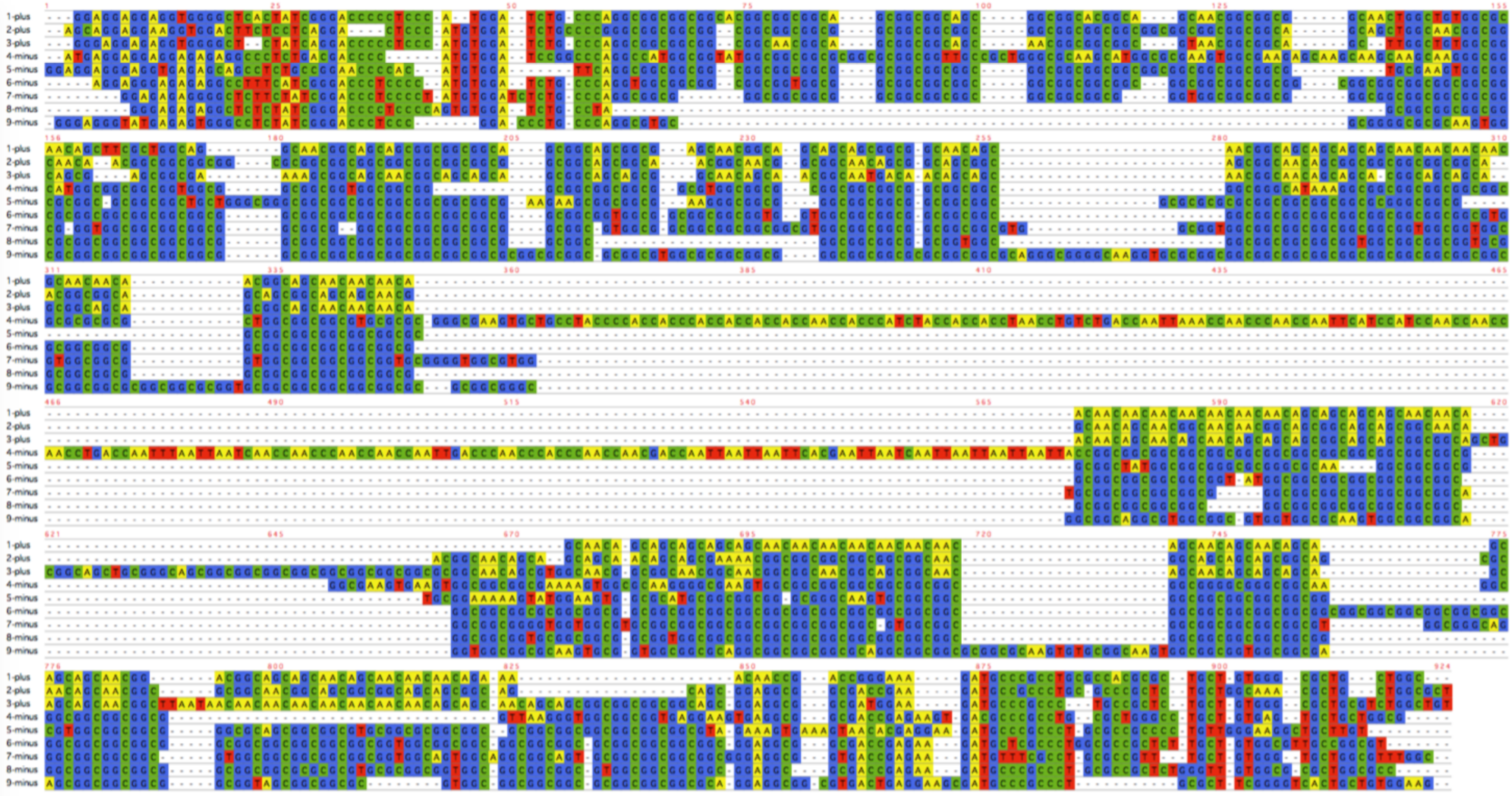
1-plus
2-plus
3-plus
4-minus
5-minus
6-minus
7-minus
8-minus
9-minus

F9-1

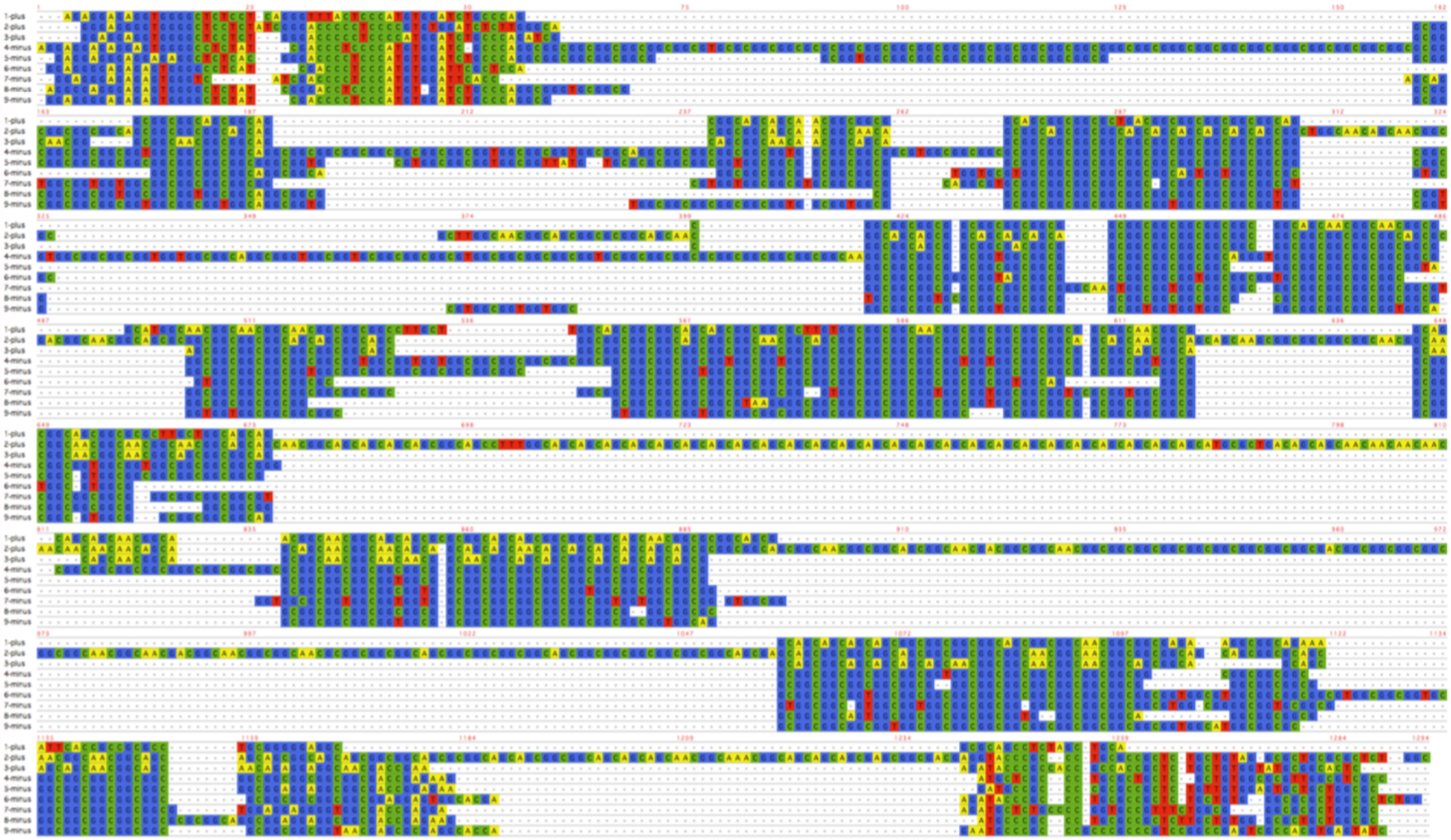

FD-1

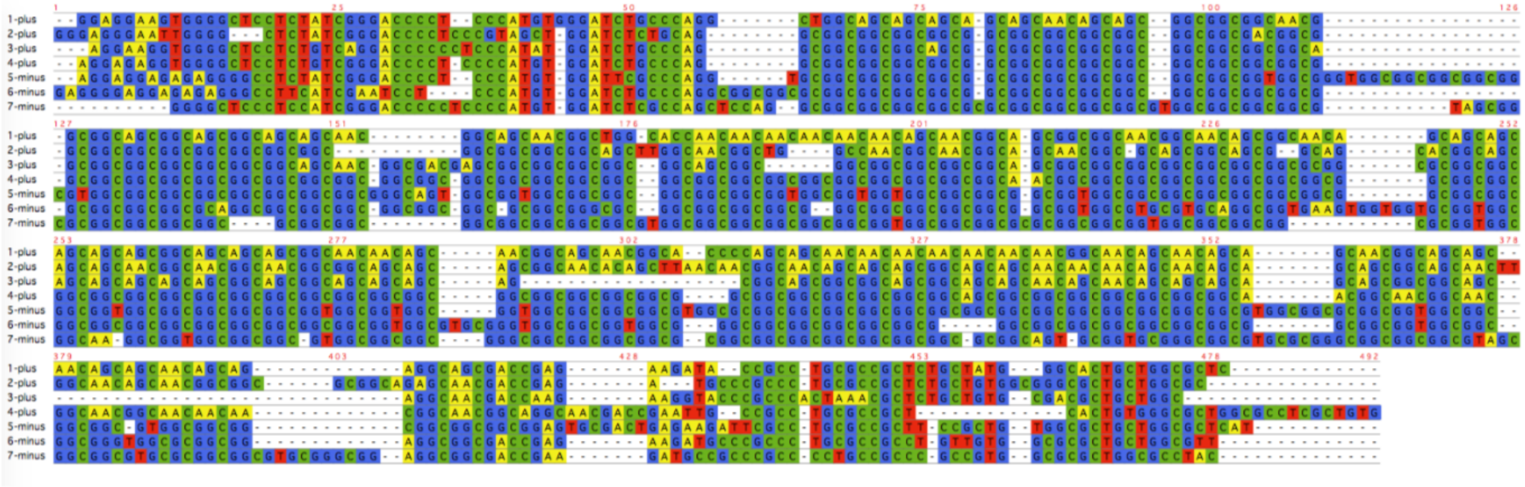

FD-2

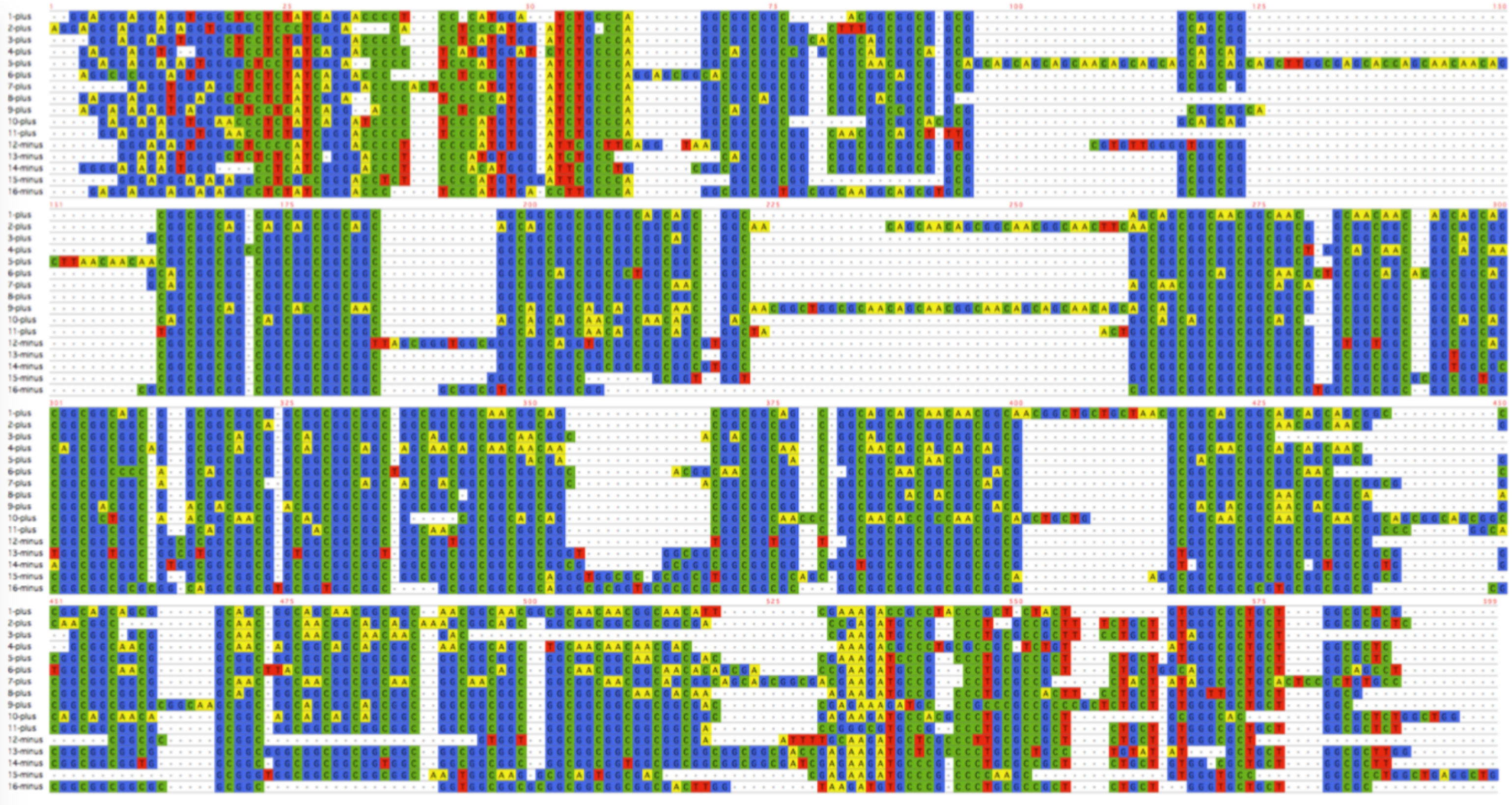

**Supplementary Figure 2. Multiple-alignment of sequences from expanded repeats in affected individuals.**

There is a possibility that the expanded repeat contains GGC and GGA repeats in patients from family1, 2 and 3. GGA repeats seem to be present towards the 3’ end of the repeat expansion. In F-1, asterisks indicate where both plus and minus strands have GGA, suggesting these might not be nanopore sequencing errors.

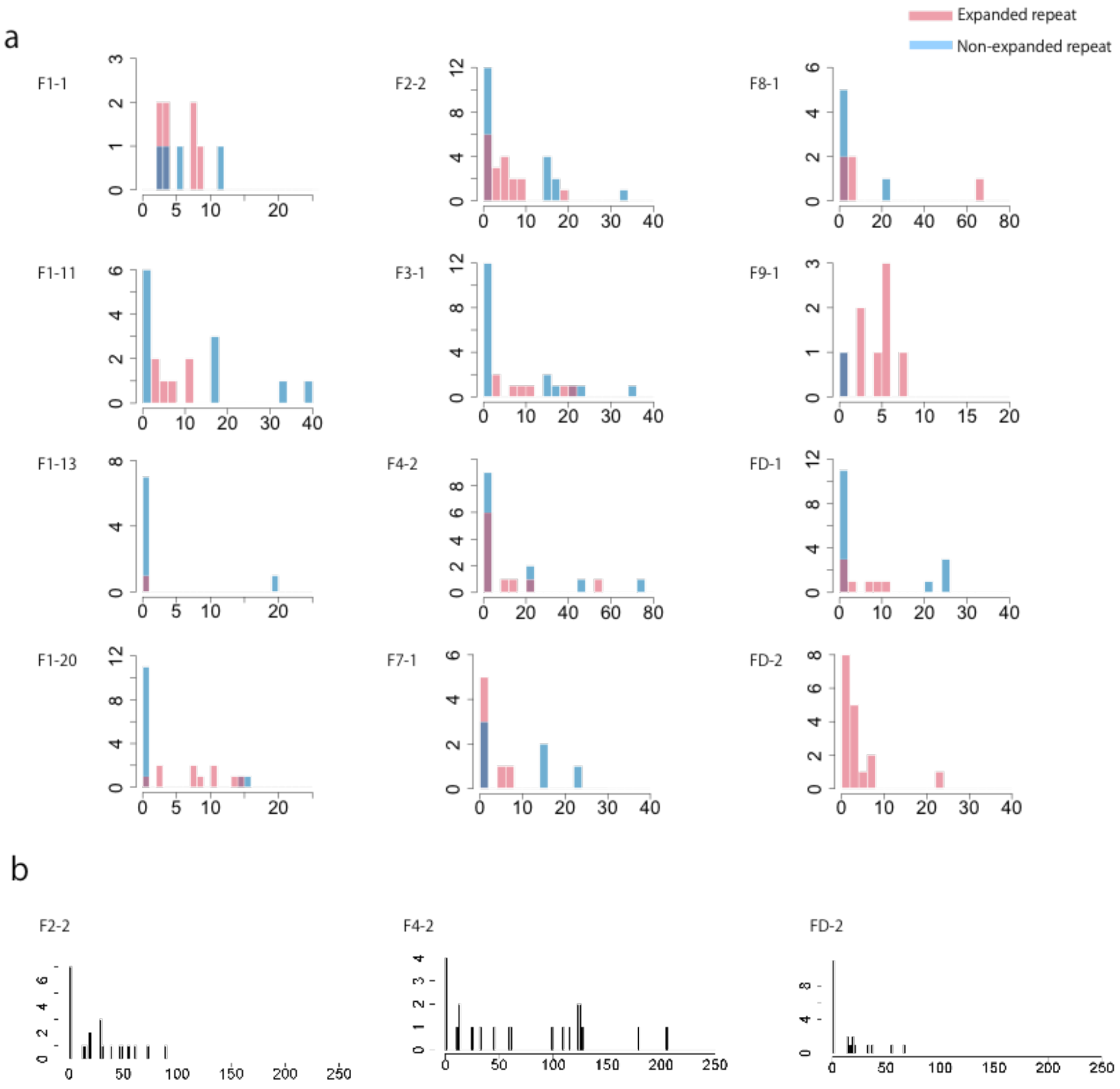


**Supplementary Figure 3. The numbers of 5mC in *NOTCH2NLC* repeat.**

**a.** The numbers of 5mC in *NOTCH2NLC* repeats (chr1:149390802-149390842) in both expanded and non-expanded repeats are shown (pale pink: expanded, pale blue: non-expanded). There is no difference between expanded and non-expanded repeats in 5mC modification of cytosine in any of the NIID patients. y-axis: read count, x-axis: number of flappie-called 5mC in 1,000 bases. **b.** Imprinted region in chr15 CpG island [12] from three individuals with largest datasets (F2-2, F4-2 and FD-2) are shown as positive controls for flappie 5mC methylation detection. This region of the maternal allele should be methylated. As expected, it shows some tendency to bimodal distribution.

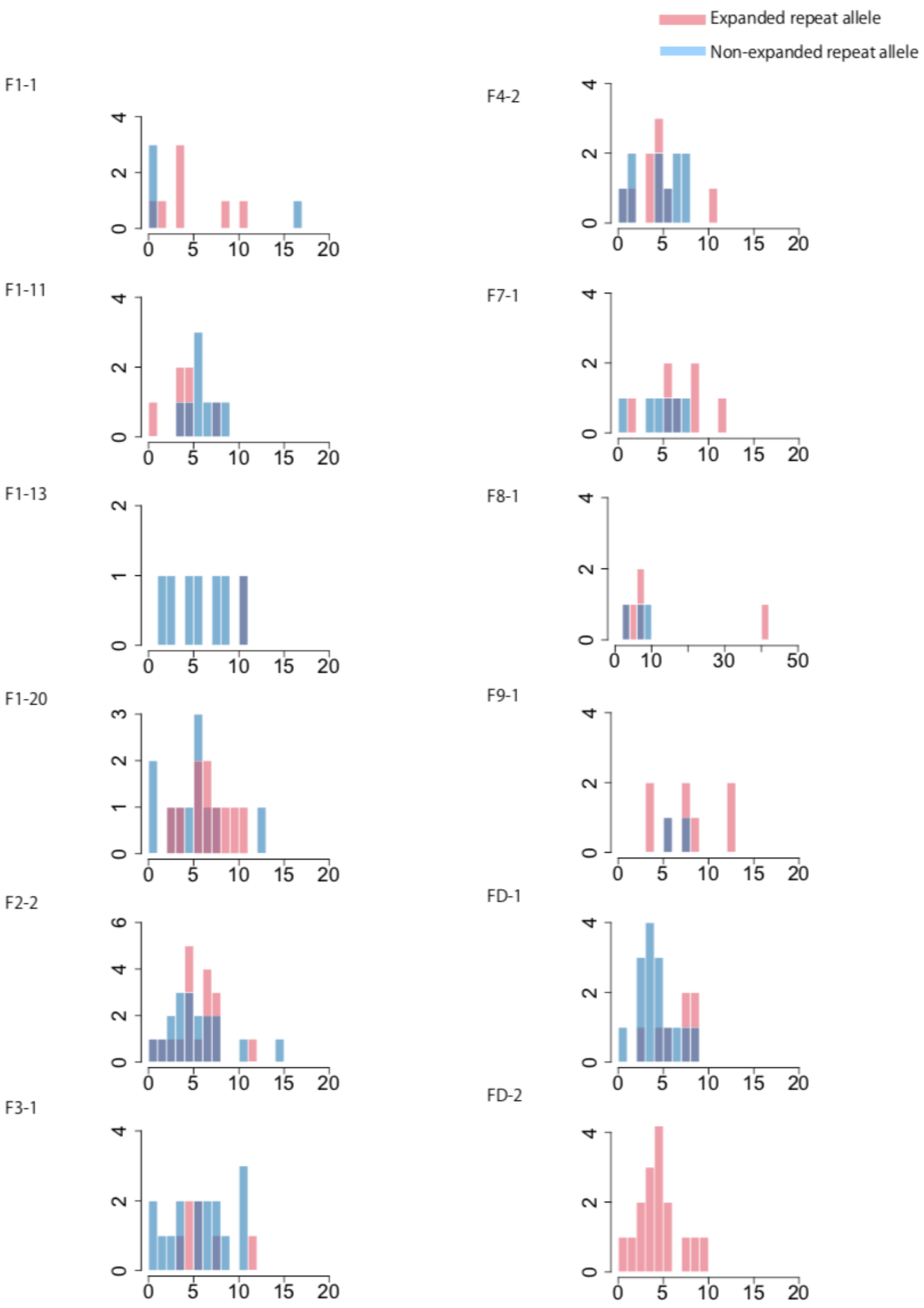


**Supplementary Figure 4. The numbers of 5mC downstream of *NOTCH2NLC* repeat.**

The numbers of 5mC modifications in a CpG island (chr1:149390845-149391541), which is downstream of the *NOTCH2NLC* repeat. The CpG island methylation of expanded and non-expanded repeat alleles (chr1:149390802-149390842) is shown separately. There is no difference between expanded and non-expanded repeat alleles in 5mC modification in any of the NIID patients. Pale pink: CpG on expanded allele, pale blue: CpG on non-expanded allele. y-axis: read count, x-axis: number of flappie-called 5mC in 1,000 bases.

### Supplementary Tables

| individual ID | disease status | sequencer | number of reads | total base | Median | average read length | predicted coverage | total number of tandem-genotypes call | number of read with repeat expantion | prioritization with single data | prioritization with 4 control data |
|---|---|---|---|---|---|---|---|---|---|---|---|
| F1-1 | affected | PromethION | 2,112,375 | 19,006,257,320 | 3520 | 8,998 | 6 | 9 | 7 | 1 | 1 |
| F1-3 | unaffected | PromethION | 23,303,818 | 54,535,886,940 | 691 | 2,340 | 17 | 11 | 0 | NA | NA |
| F1-6 | affected | PacBio RSII | 9,064,262 | 74,975,739,315 | 7701 | 8,272 | 23 | 14 | 8 | 1 | 1 |
| F1-9 | unaffected | PromethION | 9,635,261 | 35,004,315,947 | 2452 | 3,633 | 11 | 19 | 0 | NA | NA |
| F1-11 | affected | PromethION | 10,589,493 | 57,885,853,963 | 3298 | 5,466 | 18 | 13 | 6 | 3 | 1 |
| F1-12 | unaffected | PromethION | 9,091,759 | 63,874,106,363 | 5889 | 7,026 | 19 | 19 | 0 | NA | NA |
| F1-13 | affected | PromethION | 4,790,029 | 27,954,971,034 | 4364 | 5,836 | 8 | 9 | 2 | 3 | 1 |
| F1-20 | affected | PromethION | 12,830,261 | 57,451,863,375 | 1558 | 4,478 | 17 | 22 | 11 | 3 | 1 |
| F2-2 | affected | PromethION | 15,926,839 | 82,870,233,437 | 3520 | 5,203 | 25 | 32 | 16 | 2 | 1 |
| F2-3 | unaffected | PromethION | 18,339,946 | 23,268,659,841 | 865 | 1,269 | 7 | 6 | 0 | NA | NA |
| F3-1 | affected | PromethION | 11,109,622 | 62,195,375,599 | 2990 | 5,598 | 19 | 21 | 6 | 2 | 2 |
| F4-2 | affected | PromethION | 10,658,181 | 72,698,332,750 | 3736 | 6,821 | 22 | 22 | 13 | 1 | 1 |
| F7-1 | affected | PromethION | 5,529,417 | 38,773,128,508 | 3169 | 7,012 | 12 | 13 | 7 | 1 | 1 |
| F8-1 | affected | PromethION | 17,969,232 | 49,838,565,927 | 1608 | 2,774 | 15 | 11 | 9 | 2 | 1 |
| F9-1 | affected | PromethION | 6,786,707 | 55,398,873,781 | 3520 | 8,163 | 17 | 13 | 9 | 1 | 1 |
| FD-1 | affected | PromethION | 11,467,114 | 56,214,013,607 | 3110 | 4,902 | 17 | 21 | 7 | 5 | 5 |
| FD-2 | affected | PromethION | 13,655,084 | 78,995,795,042 | 3861 | 5,785 | 24 | 16 | 16 | 5 | 3 |

**Supplementary Table 1.** Summary of dataset information and tandem-genotypes results. Prioritization ranks of tandem-genotypes for the *NOTCH2NLC* repeat out of all the one million tandem repeats throughout the whole genome are also shown. Four control datasets were used for prioritization are data 3, 19 (public data) and F1-3, -9 (unaffected family members).

| data | sequencer | total number of tandem-genotypes call at NOTCH2NLC repeat | number of reads | total base | average length |
|---|---|---|---|---|---|
| 1 | MinION | 2 | 2,239,370 | 18,159,653,811 | 8,109 |
| 2 | MinION | 18 | 5,246,726 | 45,134,820,953 | 8,603 |
| 3 | MinION | 27 | 14,183,584 | 91,240,120,433 | 6,433 |
| 4 | PromethION | 21 | 6,003,887 | 46,621,512,542 | 7,765 |
| 5 | PromethION | 25 | 3,394,312 | 43,196,945,412 | 12,726 |
| 6 | PromethION | 28 | 5,074,319 | 96,709,785,254 | 19,059 |
| 7 | PromethION | 28 | 4,497,556 | 90,866,959,081 | 20,204 |
| 8 | PromethION | 24 | 4,403,236 | 81,554,440,718 | 18,522 |
| 9 | PromethION | 32 | 8,642,604 | 111,990,048,861 | 12,958 |
| 10 | PromethION | 19 | 9,635,261 | 35,004,315,947 | 3,633 |
| 11 | PromethION | 30 | 7,136,355 | 86,634,317,185 | 12,140 |
| 12 | PromethION | 11 | 23,303,818 | 54,535,886,940 | 2,340 |
| 13 | PromethION | 19 | 7,824,636 | 77,878,925,289 | 9,953 |
| 14 | PromethION | 15 | 3,557,359 | 53,696,135,713 | 15,094 |
| 15 | PromethION | 8 | 1,885,376 | 19,822,791,751 | 10,514 |
| 16 | PromethION | 7 | 2,981,694 | 22,136,031,084 | 7,424 |
| 17 | PromethION | 19 | 9,091,759 | 63,874,106,363 | 7,026 |
| 18 | PromethION | 6 | 18,339,946 | 23,268,659,841 | 1,269 |
| 19 | PromethION | 34 | 13,134,890 | 185,056,318,464 | 14,089 |
| 20 | Pacbio Sequel | 19 | 6,174,383 | 44,835,599,221 | 7,262 |
| 21 | Pacbio Sequel | 9 | 4,029,436 | 39,348,377,910 | 9,765 |
| 22 | Pacbio Sequel | 10 | 3,314,492 | 30,665,693,222 | 9,252 |
| 23 | Pacbio Sequel | 5 | 2,218,415 | 21,275,114,124 | 9,590 |
| 24 | Pacbio Sequel | 4 | 2,874,747 | 25,207,588,869 | 8,769 |
| 25 | Pacbio Sequel | 6 | 1,929,367 | 20,280,159,023 | 10,511 |
| 26 | Pacbio Sequel | 6 | 2,705,416 | 25,484,894,848 | 9,420 |
| 27 | Pacbio Sequel | 6 | 8,391,859 | 56,100,283,182 | 6,685 |
| 28 | Pacbio Sequel | 10 | 3,228,957 | 30,151,182,772 | 9,338 |
| 29 | Pacbio Sequel | 7 | 3,650,569 | 30,427,397,011 | 8,335 |

**Supplementary Table 2**. Sequencing summary of control datasets. Data 3 and 19 are publicly available, rel3 [6] and ERR2585112-5 [7], respectively.

| Family No. | Individual No. | Disease status | Long read seq. | RP-PCR |
|---|---|---|---|---|
| 1 | 1 | Affected | Expansion | Expansion |
| 1 | 2 | Unaffected | N.P. | No expansion |
| 1 | 3 | Unaffected | No expansion | No expansion |
| 1 | 6 | Affected | Expansion | Expansion |
| 1 | 7 | Unaffected | N.P. | No expansion |
| 1 | 8 | Affected | N.P. | Expansion |
| 1 | 9 | Unaffected | No expansion | No expansion |
| 1 | 10 | Affected | N.P. | Expansion |
| 1 | 11 | Affected | Expansion | Expansion |
| 1 | 12 | Unaffected | No expansion | No expansion |
| 1 | 13 | Affected | Expansion | Expansion |
| 1 | 14 | Affected | N.P. | Expansion |
| 1 | 16 | Affected | N.P. | Expansion |
| 1 | 17 | Unaffected | N.P. | No expansion |
| 1 | 18 | Unaffected | N.P. | No expansion |
| 1 | 20 | Affected | Expansion | Expansion |
| 2 | 1 | Unaffected | N.P. | No expansion |
| 2 | 2 | Affected | Expansion | Expansion |
| 2 | 3 | Unaffected | No expansion | No expansion |
| 3 | 1 | Affected | Expansion | Expansion |
| 4 | 1 | Affected | N.P. | Expansion |
| 4 | 2 | Affected | Expansion | Expansion |
| 4 | 3 | Affected | N.P. | Expansion |
| 4 | 4 | Affected | N.P. | Expansion |
| 7 | 1 | Affected | Expansion | Expansion |
| 7 | 2 | Affected | N.P. | Expansion |
| 8 | 1 | Affected | Expansion | Expansion |
| 8 | 2 | Affected | N.P. | Expansion |
| 9 | 1 | Affected | Expansion | Expansion |
| 9 | 2 | Affected | N.P. | Expansion |
| 10 | 1 | Affected | N.P. | Expansion |
| D | 1 | Affected | Expansion | Expansion |
| D | 2 | Affected | Expansion | Expansion |

**Supplementary Table 3. Comparison of long read sequencing and RP-PCR results in 9 NIID families.**

N.P. Not performed.

| Individual No. | Disease status | Long read seq. | RP-PCR |
|---|---|---|---|
| 3614 | Affected | N.P. | Expansion |
| 3615 | Affected | N.P. | Expansion |
| 3616 | Affected | N.P. | Expansion |
| 3618 | Affected | N.P. | Expansion |
| 3619 | Affected | N.P. | Expansion |
| 3623 | Affected | N.P. | Expansion |
| 3624 | Affected | N.P. | Expansion |
| 3625 | Affected | N.P. | Expansion |
| 3626 | Affected | N.P. | Expansion |
| 3627 | Affected | N.P. | Expansion |
| 3628 | Affected | N.P. | Expansion |
| 3630 | Affected | N.P. | Expansion |
| 3636 | Affected | N.P. | Expansion |
| 3639 | Affected | N.P. | Expansion |
| 3644 | Affected | N.P. | Expansion |
| 3645 | Affected | N.P. | Expansion |
| 3646 | Affected | N.P. | Expansion |
| 3647 | Affected | N.P. | Expansion |
| 3648 | Affected | N.P. | Expansion |
| 3650 | Affected | N.P. | Expansion |
| 3651 | Affected | N.P. | Expansion |
| 3652 | Affected | N.P. | Expansion |
| 3653 | Affected | N.P. | Expansion |
| 3654 | Affected | N.P. | Expansion |
| 3658 | Affected | N.P. | Expansion |
| 3661 | Affected | N.P. | Expansion |
| 3662 | Affected | N.P. | Expansion |
| 3663 | Affected | N.P. | Expansion |
| 3665 | Affected | N.P. | Expansion |
| 3667 | Affected | N.P. | Expansion |
| 3668 | Affected | N.P. | Expansion |
| 3672 | Affected | N.P. | Expansion |
| 3673 | Affected | N.P. | Expansion |
| 3676 | Affected | N.P. | Expansion |
| 3677 | Affected | N.P. | Expansion |
| 3680 | Affected | N.P. | Expansion |
| 3683 | Affected | N.P. | Expansion |
| 3687 | Affected | N.P. | Expansion |
| S81 | Affected | N.P. | Expansion |

**Supplementary Table 4. RP-PCR results in 39 sporadic NIID cases.**
N.P. Not performed.
